# A method for single molecule localization microscopy of tissues reveals non-random distribution of nuclear pores in *Drosophila*

**DOI:** 10.1101/2021.05.24.445468

**Authors:** Jinmei Cheng, Edward S. Allgeyer, Jennifer H. Richens, Edo Džafić, Amandine Palandri, Bohdan Lewkow, George Sirinakis, Daniel St Johnston

**Affiliations:** The Gurdon Institute & the Department of Genetics, University of Cambridge, Tennis Court Road, Cambridge, CB2 1QN, UK; Institute of Reproductive Medicine, School of Medicine, Nantong University, Nantong 226001, China; Institute of Molecular Health Sciences, ETH Zurich, Zurich, Switzerland

## Abstract

Single Molecule Localisation Microscopy (SMLM) can provide nanoscale resolution in thin samples but has rarely been applied to tissues, because of high background from out of focus emitters. Here we describe a line scanning microscope that provides optical sectioning for SMLM in tissues. Imaging endogenously-tagged nucleoporins and F-actin on this system using DNA- and peptide-PAINT routinely gives 30nm resolution or better at depths greater than 20 µm. This revealed that the nuclear pores are nonrandomly distributed in most *Drosophila* tissues, in contrast to cultured cells. Lamin Dm_0_ shows a complementary localisation to the nuclear pores, suggesting that it corrals the pores. Furthermore, ectopic expression of the tissue-specific Lamin C distributes the nuclear pores more randomly, whereas *lamin C* mutants enhance nuclear pore clustering, particularly in muscle nuclei. Since nucleoporins interact with specific chromatin domains, nuclear pore clustering could regulate chromatin organisation locally and contribute to the disease phenotypes caused by human Lamin A/C laminopathies.

## Introduction

The development of super-resolution microscopy techniques that break the diffraction limit has enabled the visualisation of subcellular structures with unprecedented resolution^1,2^. Some of the highest resolution is provided by a range of Single Molecule Localization Microscopy (SMLM) approaches that separate the fluorescence produced by closely-spaced fluorophores in time, so that each camera frame captures the image of only a few sparsely distributed, active emitters ^3-5^. The image of each emitter is fitted with a model function, typically a 2-dimensional (2D) Gaussian, to localize the true emitter position within some uncertainty bound. Repetitively localizing thousands or millions of single emitter positions over many camera frames results in a reconstructed final image with a resolution below the diffraction limit, typically on the order of 20 to 30 nm.

The localization precision, and subsequently the resolution, achieved by SMLM depends on the number of photons from each blink and the background, primarily from out of focus emitters. However, this has largely constrained SMLM to thin samples close to the coverslip, where the background is low. The resolution can be further enhanced by reducing the background with illumination configurations that restrict the excitation of the fluorophores to a narrow axial region, such as total internal reflection fluorescence (TIRF), highly inclined and laminated optical sheet illumination (HILO) or selective plane illumination (SPIM)^6-8^ .

SMLM imaging of thick tissues is more challenging, because of the high levels of background fluorescence from out of focus emitters and tissue autofluorescence, as well as the reduced number of photons collected from each emission due to aberrations or scattering. Chemical fluorophores have higher stability and quantum yields than fluorescent proteins, making SMLM techniques that are based on fluorescent dyes better suited for tissue samples. In the most common SMLM technique, STORM, specific “blinking” dyes are conjugated to the protein of interest, switched into a dark state and then imaged as they randomly reactivate in a special “blinking buffer” ^5,9^. The range of dyes that can be induced to blink is limited, however, and each dye has its own optimal buffer and blinking characteristics, which complicates multicolour imaging. Photobleaching can also reduce image quality as fluorophores that bleach before they are imaged are not detected ^10^. These problems are largely avoided by an alternative labelling approach called DNA-based point accumulation for imaging in nanoscale topography (DNA-PAINT), in which the protein of interest is labelled with a docking oligonucleotide (the docking strand) and then imaged using the complementary oligonucleotide (the imager strand) fused to a fluorescent probe^11^. The imager strand diffuses rapidly in solution and appears as background, but remains stationary on hybridising to the docking strand, producing a “blink” of light that can then be localised. Because the fluorophore is not directly conjugated to the protein of interest and there is a large excess of imager strand in solution, the sample does not suffer from photobleaching. One can therefore image for longer and collect many “blinks” from the same labelled protein. DNA-PAINT also removes the need to use fluorophores that “blink” in specialized buffers, and one can therefore more flexibly choose pairs of spectrally-separated fluorophores with high photon yields, improving two colour imaging. On the other hand, DNA-PAINT has higher background due to fluorescence from freely diffusing imager strands.

One imaging modality that improves imaging performance in thick samples is confocal microscopy which relies on a mechanical pinhole to reject out of focus light. Although confocal microscopy is most often realized in a point scanning geometry, it is unsuitable for SMLM imaging because it is intrinsically slow. However, camera-based variants such as spinning disk or line scanning confocal microscopes have recently been employed in SMLM to image the entire thickness of cells and, in some cases, tissue ^12,13^. However, even with these developments, SMLM is generally limited to cultured cell applications, as the localization precision deteriorates with imaging depth in high-background environments.

Here we develop a confocal line scanning system which utilizes the rolling shutter capabilities of a commercially available scientific complementary metal–oxide– semiconductor (sCMOS) camera sensor and allows SMLM imaging in tissue samples, or other high background environments such as DNA-PAINT. We present a pipeline for performing multi-colour SMLM on endogenous proteins in tissue samples, using optimised protocols for labelling SNAP and Halo-tagged proteins with docking strand oligonucleotides and a modified version of the IRIS probe for visualising F-actin as a landmark. We tested the line scanning system using endogenously-tagged nucleoporins and discovered that the nuclear pores are not randomly distributed in most *Drosophila* tissues, but are corralled into specific regions of the nuclear envelope by a Lamin-dependent mechanism.

## Results

### Design of a microscope for PAINT imaging in tissues

To perform optical sectioning of thick samples, we designed and built an inexpensive line scanning confocal system using standard optical components (**Fig. 1a** and **Supplementary Fig. 1**).

**Figure 1.**
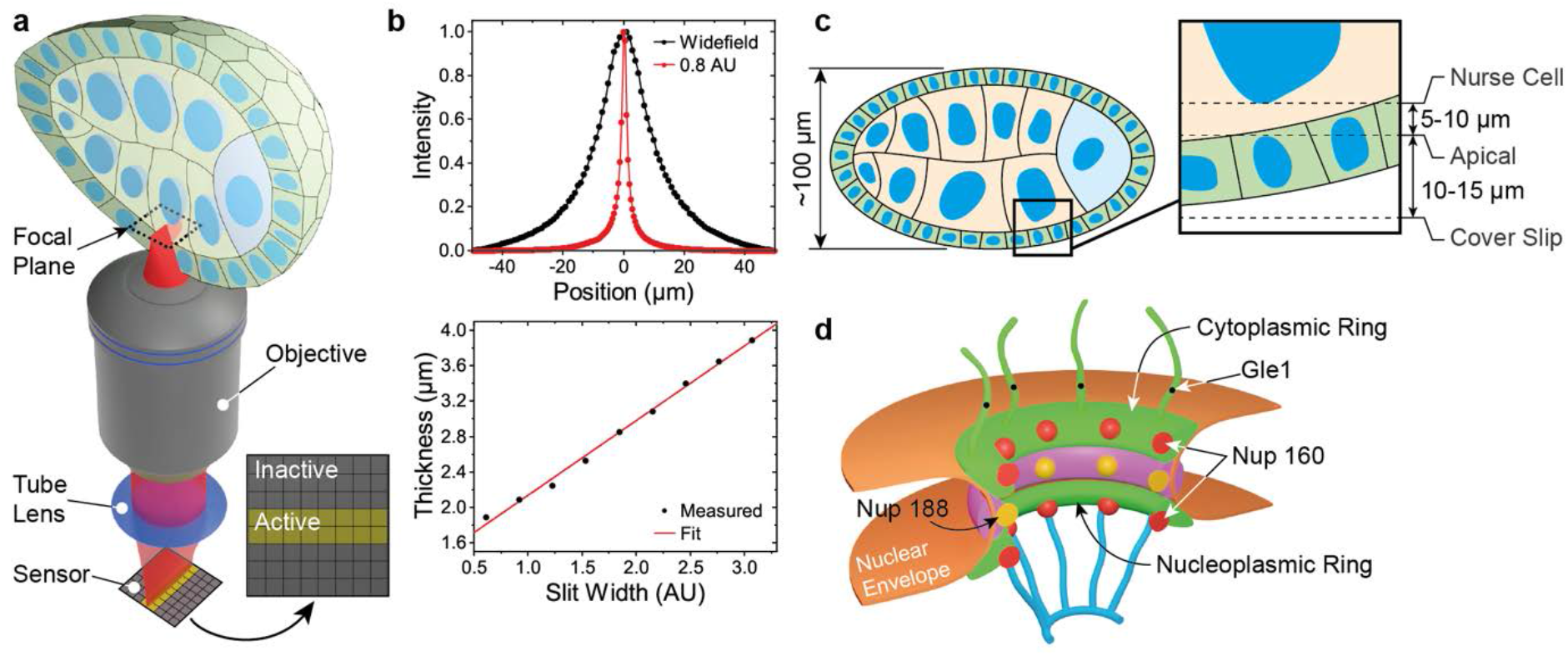
Microscope and sample geometry overview. **a** Excitation light is focused into a line at the sample plane inside the egg chamber. Fluorescence is collected by the objective lens and replayed to an sCMOS camera chip where active and inactive pixel rows form an electronic slit for optical sectioning. **b** Optical thickness comparison between widefield (normal) and slit scanning modes. An optical section of ∼1.5 µm is formed with the electronic slit set to 0.8 airy units. **c** Detailed view of a typical image focal plane. The apical membrane resides 10-15 µm inside the egg chamber while the nurse cell nucleus is an additional 5-10 µm further depending on egg chamber stage. **d** Schematic of a nuclear pore complex. Nup160 resides in the outer cytoplasmic and nucleoplasmic rings, Nup188 lies in the inner rings and Gle1 is a component of the cytoplasmic filaments.

Briefly, an excitation laser is focused into a line with diffraction limited width at the sample and scanned across the field of view (FOV) with a galvanometer mirror (**Supplementary Fig. 1**). Fluorescence is collected with the same objective and relayed to an sCMOS camera. In contrast to previous implementations of confocal line scanning that employed a mechanical slit, we exploited the flexible architecture of the sCMOS camera and used rows of active and inactive pixels to form an “electronic” slit that rejects out of focus light^14^. By adjusting the time delay between neighbouring rows, one can set the speed with which the active pixel rows move across the camera sensor and synchronize it with the scanning excitation line to form an image. The width of the electronic slit can be controlled by the software and has the same effect as opening or closing a confocal pinhole to adjust the optical section thickness (**Fig. 1b**). In this way, pixel integration time, slit width, and scanning speed can be optimized to maximize signal collection and minimize out-of-focus background (**See Supplementary information**). The use of an “electronic” slit minimizes mechanical parts and allows for a simple optical setup with tuneable optical sectioning superior to wide-field illumination (**Fig. 1c**).

### Testing microscope performance by imaging NPC in tissues

To examine the performance of the microscope, we imaged *Drosophila* egg chambers (**Fig. 1c**), as these are amenable to genetic manipulation and large enough to test the optical sectioning capabilities of the system. The nuclear pore complex (NPC) has been proposed as a reference structure for super-resolution microscopy, since it is abundant in the nuclear envelopes of all cells and has a known structure and stoichiometry, with a characteristic 8-fold symmetry^15,16^. To image endogenous proteins using DNA-PAINT, we used CRISPR-mediated homologous recombination to introduce the self-labelling SNAP and Halo tags into three nucleoporins that localise to different parts of the NPC: Nup160, which is present in two copies in each of the 8 spokes of the cytoplasmic and nuclear outer rings, Nup188 found in 32 copies in the inner ring and Gle1, a component of the cytoplasmic filaments that extend from near the centre of the NPC (**Fig. 1d**)^16-18^.

To perform DNA-PAINT imaging of nuclear pores, we optimised conditions for conjugating docking strands to the tagged nucleoporins. Docking strands can be efficiently conjugated to SNAP and Halo-tagged proteins by adding O6-benzylguanine or the chloroalkane HaloTag ligand to the 5’ end of the oligonucleotide **(Supplementary Fig. 2a)**, as previously reported^19^. Promega also kindly provided us with a compound, PBI-300-43, that can be attached to a 12-Carbon linker on the 5’ end of the oligonucleotide to produce a modified docking strand that is a better substrate for the Halo Tag in some in vitro assays (**Supplementary Fig. 2b-c**). Both chloroalkane-modified oligonucleotides conjugated equally efficiently to the Halo-tagged proteins in fixed and permeabilised *Drosophila* egg chambers. We measured the effective labelling efficiency (EL) for Halo by imaging Nup160 and Gle1 in flies that were homozygous for Nup160-SNAP and heterozygous or homozygous for Gle1-Halo and scoring the proportion of Nup160-marked nuclear pores that contained Gle1-Halo signal in the centre. There are 8 copies of Gle1 per nuclear pore and the probability of any Gle1 molecule being detected in Gle1 heterozygotes is half the effective labelling efficiency because only one of the two copies are tagged. The probability of a nuclear pore containing no detected Gle1 molecules is therefore 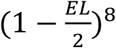^8^ for Gle1 heterozygotes and (1 − *EL*)^8^ for homozygotes allowing us to calculate EL from the frequency of nuclear pores with no Gle1 signal. These measurements gave effective labelling efficiencies of 20-40%, with fresh docking strands giving higher labelling (**Supplementary Table 1**). This is comparable to the labelling efficiencies obtained when conjugating nucleotides to SNAP-tagged proteins^19^.

**Figure 2.**
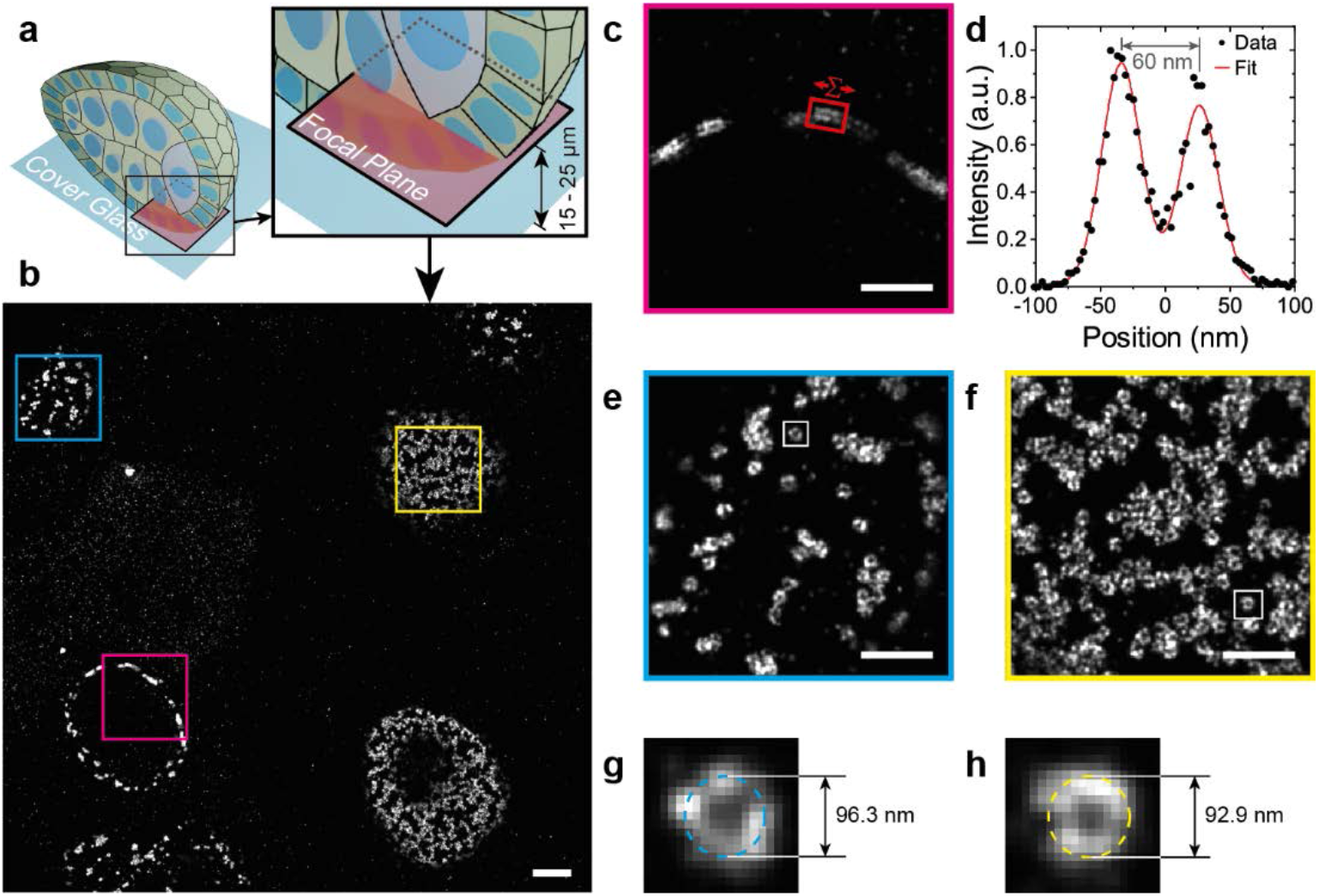
DNA-PAINT Egg Chamber Imaging. **a** An Illustration of the position of the focal plane imaged in (**b**), which is 15 to 25 µm above the coverslip and inside the egg chamber. **b** DNA-PAINT image showing nurse cell and follicle cell nuclei. **c** A magnified view of the area outlined by the red box in **b** showing a cross-sectional view of NPCs in the nuclear membrane. The intensity distribution between the outer and inner rings of the NPCs in region in **c** is shown in **d**, with the distance measured as 60 nm. Detailed views of NPCs in the follicle cell nuclear membrane (**e**) and the nurse cell nuclear membrane (**f**), showing distinct ring structures with hollow centres. **g** and **h** show detailed views of NPCs from **e** and **f** fit with a circle and diameters of 96.3 and 92.9 respectively.

We used this method to perform DNA-PAINT SMLM of the nuclear pores in the follicle cells (**Fig. 2b, and e**) and germ cells (**Fig. 2b and 2f**) of *Drosophila* egg chambers homozygous for Nup160-Halo. Since DNA-PAINT is resistant to photobleaching, we could routinely collect 50,000 to 100,000 camera frames with only a 20 – 40% reduction in the number of blinks per frame by the end (see **Supplementary Figure 3a**), presumably due to photodamage to the docking strands when bound imager strands are bleached^20^. We obtained between 550 and 2200 photons per blink, depending on the imaging conditions (see **Supplementary Figure 3b**), to give a localisation precision between 7 and 12 nm (see **Supplementary Figure 3c**), resulting in a maximum achievable resolution of 17 to 29 nm. This resolution allowed us to measure the average distance between the nuclear and cytoplasmic rings of the NPC in our images as 58.7 ± 4.4 nm (mean value ± standard deviation, n = 99, **Supplementary Fig. 4**), which corresponds well with the distance measured from EM data^21^ (**Fig. 2d**). The mean NPC diameter was also measured by performing a circular fit on individual NPCs (**Fig. 2g, h**) and found to be 103.4 ± 6.5 nm (mean value ± standard deviation, n = 226, **Supplemental Fig. 5**) in good agreement with the expected value from EM on *Drosophila* NPCs and super-resolution studies of NPCs in other organisms^15,22^. The nurse cell nuclei lie approximately 10 µm above the follicle cell epithelium, which is at least 5 µm thick (**Fig. 2a**). Thus, the imaging plane in **Fig. 2** is ≥ 15 µm into the sample, demonstrating that our line scanning system can collect high quality super-resolution images deep inside tissues.

**Figure 3.**
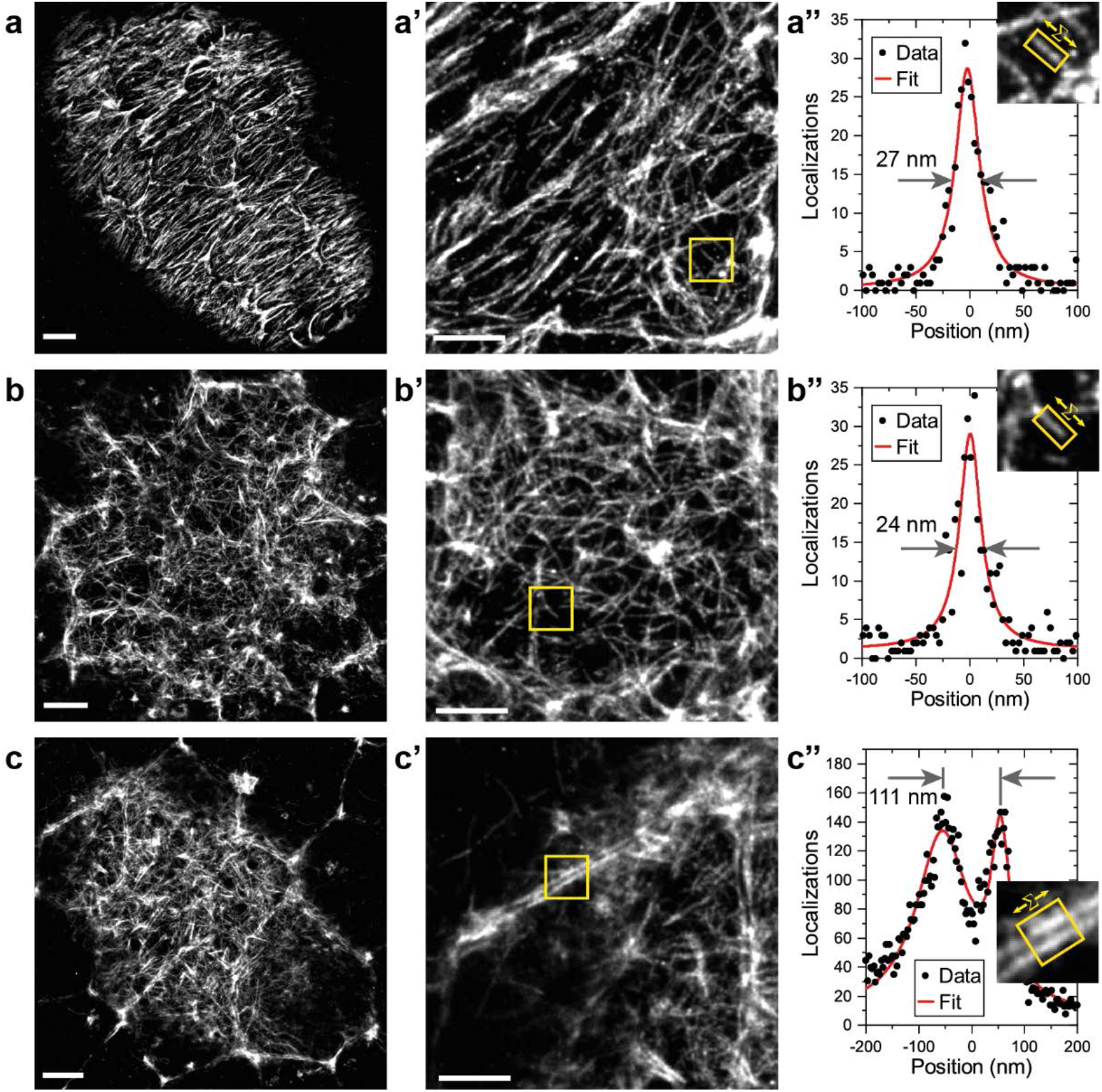
Peptide-PAINT Imaging in *Drosophila* Egg Chambers. **a** Basal F-actin in a stage 5 *Drosophila* egg chamber. **a’** Detailed view from **a** showing that smaller filaments are well-resolved. **a’’** A single filament from the boxed region in **a’** is shown on the upper right. The graph shows the distribution of localizations perpendicular to the straight filament boxed in yellow after summing along its long axis (see **Supplementary Methods** for details). The filament diameter was measured as 27 nm FWHM by fitting with a Gaussian. **b** Apical F-actin in a stage 5 *Drosophila* egg chamber. **b’** Detailed view from **b** showing small filaments are visible. **b’’** The box region in **b’** is shown at upper right. The graph shows the distribution of localizations perpendicular to the straight filament boxed in yellow after summing along its long axis. The filament diameter was measured as 24 nm FWHM by fitting with a Gaussian. **c** Apical F-actin in a stage 8 *Drosophila* egg chamber. **c’** Detailed view from **c** showing the clear separation between the cortical F-actin on adjacent cell membranes. **c’’** The boxed region in **c’** is shown on the upper right. The graph shows the distribution of blinks perpendicular to the regions of the cell cortices boxed in yellow after summing the localizations along each cell cortex. Fitting with two Gaussians gives the distance between the membranes as 111 nm. Scale bars are 2 µm for panels a-c and 1 µm for panels a’-c’.

**Figure 4.**
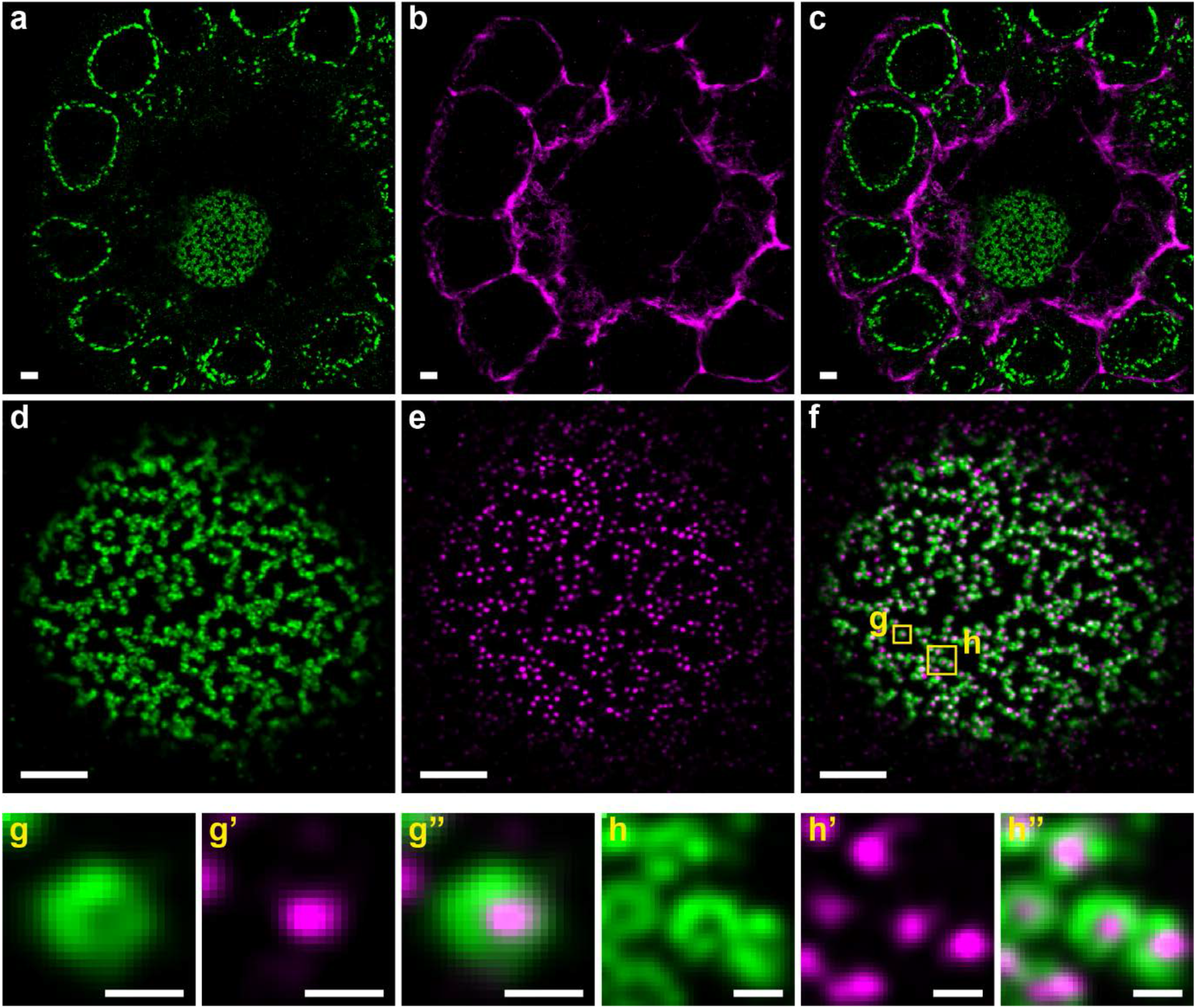
Combined DNA- and Peptide-PAINT Two Colour Imaging. **a** Nup160 visualized using DNA-PAINT and imaged simultaneously with (**b**) F-actin visualized with Peptide-PAINT. **c** Merge of **a** and **b. d** Nup160 and (**e**) Gle1 imaged simultaneously with DNA-PAINT. The merged image is shown in (**f**). **g-g”** An enlarged view of the left-hand boxed region in (**f**) showing the Nup160 signal (**g**), the Gle1 signal (**g’)** and the merged image (**g’’). h-h”**. A similar enlarged view of the right-hand boxed region in **(f)**. Gle1 appears inside the Nup160 rings as expected. Scale bars are 1 µm for panels a-f and 100 nm for g and h.

**Figure 5.**
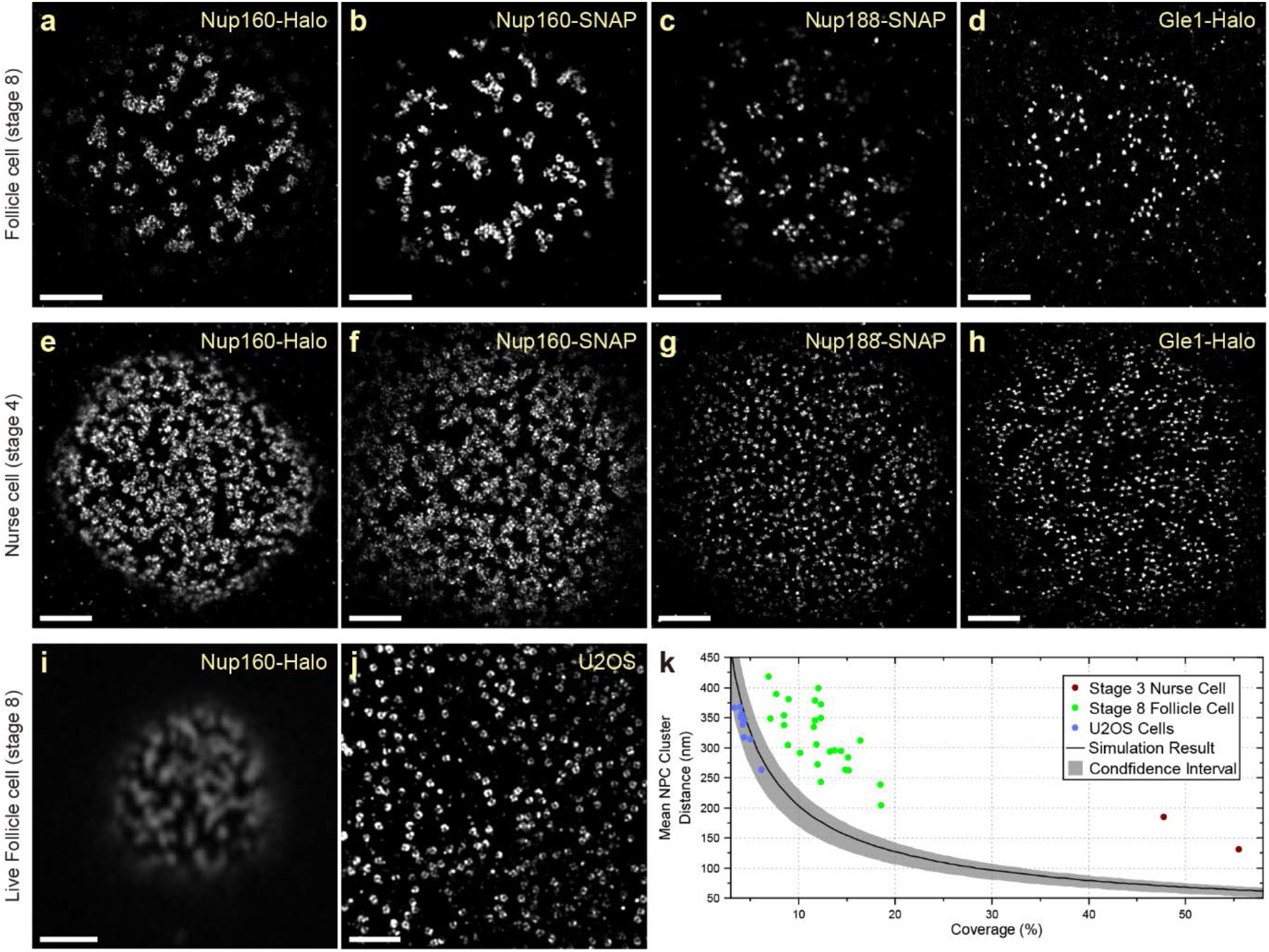
Super-Resolution Imaging in Tissue Reveals a Nonrandom Distribution of NPCs. **a-d** DNA-PAINT images of Nup160-Halo **(a)**, Nup160-SNAP **(b)**, Nup188-SNAP **(c)** and Gle1-Halo **(d)** in NPC in stage 8 follicle cell nuclei. The NPC appear clustered, regardless of the protein/tag combination used, showing that the organization is not caused by the tag. **e-h** DNA-PAINT images of Nup160-Halo **(e)**, Nup160-SNAP **(f)**, Nup188-SNAP **(g)** and Gle1-Halo **(h)** in NPCs of stage 4 nurse cell nuclei. A similar NPC clustering is seen in nurse cell nuclei, suggesting that this organization is not specific to the follicular epithelium. **i** Live imaging of JF646-labelled Nup160-Halo in a stage 8 follicle cell nucleus using a Zeiss Airyscan microscope. The clustering is present in vivo, indicating that it is not caused by fixation. **j** DNA-PAINT image of Nup96-Halo in a cultured U2OS cell nucleus. The NPCs appear uniformly distributed without any underlying organization. **k** A graph showing the mean NPC intercluster distance as a function of the proportion of the nuclear envelope covered by NPCs (Coverage). The black line shows the mean distances obtained from simulations of random NPC distributions, with 95% confidence intervals shown in grey. U2OS cells show a random distribution of NPCs, whereas the mean distances between NPC clusters are significantly larger in follicle cell and nurse cell nuclei, indicating that the NPCs are non-randomly distributed. See **Supplementary Methods** for analysis details. Scale bars are 1 µm for a-h and j and 5 µm for i.

### Imaging F-actin using Lifeact peptide-PAINT

When imaging less well-characterised proteins that do not localise to known structures, it is important to have landmarks to indicate where in the cell the protein is localised. The actin cytoskeleton represents an excellent landmark, as it outlines the cell cortex and forms specific structures in different regions of the cell. We therefore adapted the IRIS method for labelling F-actin using the Lifeact peptide, which binds to F-actin with high specificity, but low affinity^23^. To increase the stability of the probe, we added an extra Cysteine to the N-terminus of the peptide and conjugated the organic fluorescent dye, Cy3B, to this residue through a stable thioether bond using Cy3B-maleimide^24^. In this peptide-PAINT approach, transient binding of the Cy3B-Lifeact peptide to F-actin results in blinks that are imaged in the same way as DNA-PAINT. We imaged the actin organization in both the basal and apical side of follicles cells in stage 5 *Drosophila* egg chambers (**Fig. 3**). Stage 5 egg chambers rotate around their anterior-posterior axes due the coordinated circular migration of the follicle cells, driven by planar polarised lamellipodia and stress fibres on their basal surfaces ^25,26^. Super-resolution Lifeact imaging of the basal surface showed the planar polarised stress fibres with thinner actin filaments branching out from them, while the high density of actin in the lamellipodia appeared as bright regions in the image (**Fig. 3a, a’**). The thinner filaments have an apparent diameter of **30**.**4 ± 8**.**42** nm (mean ± standard deviation, N = 11, see **Supplementary Figures 6 and 7**) with the smallest features being below 20 nm and offer a good indication of the resolving power of the microscope (**Fig. 3a”**). By contrast, the actin on the apical surface of stage 5 follicle cells forms a branched network, with densely labelled nodes that may correspond to Myosin puncta^27^ (**Fig. 3b, b’**). The apparent diameter of the filaments was **30**.**2 ± 4**.**48** nm (mean ± standard deviation, N = 10), with the smallest features being below 20 nm, which was comparable to that on the basal side, demonstrating that the performance of the microscope is not significantly affected over a depth of ∼10 µm. By stage 8 of oogenesis, the apical actin network has become denser, with higher accumulations of actin at the tri-cellular junctions (**Fig. 3c, c’**). The higher resolution also allowed us to resolve the actin cortices in adjacent cells, which are separated by 111.0 ± 15.7 nm (mean ± standard deviation, N = 11, **Fig. 3c”** and **Supplementary Figure 8**).

**Figure 6.**
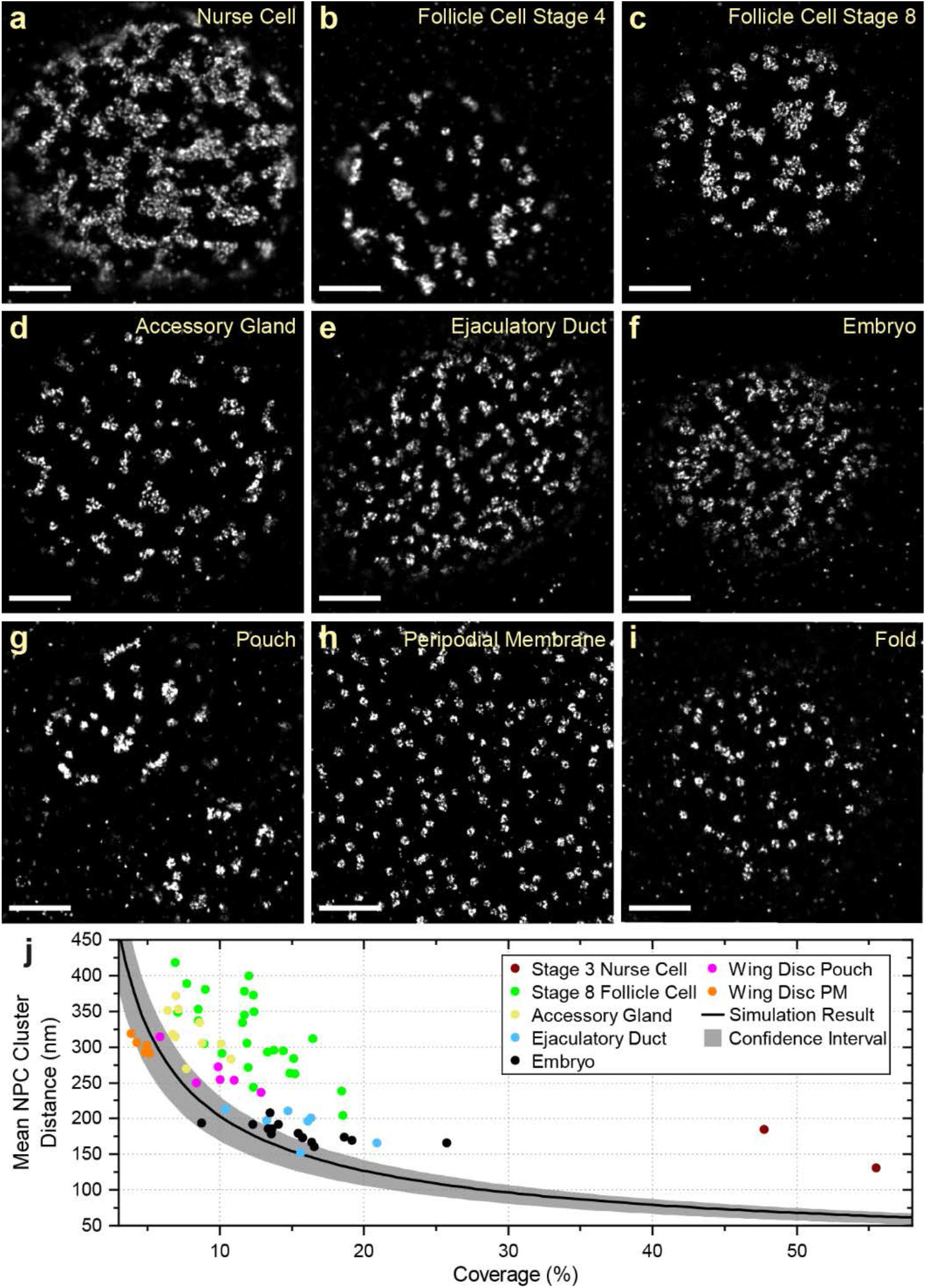
NPC organization in different *Drosophila* tissues. NPC clustering is observed in nurse cells (**a**) and follicle cells at multiple stages (**b-c**). Similar NPC clustering is observed in the accessory gland (**d**), the ejaculatory duct (**e**), the syncytial blastoderm embryo (**f**) and the pouch region of the wing imaginal disc (**g**). This organization is not observed in the peripodial membrane of the wing disc (**h**) or the wing disc fold (**i**). **j** A graph showing the mean NPC cluster distance as a function of the proportion of the nuclear envelope covered by NPCs for each tissue. The peripodial membrane (Wing Disc PM) has a similar mean inter NPC cluster distance to the simulations of random NPC distributions, whereas all other tissues show significantly greater inter-cluster distances, indicating a clustered distribution. The wing disc fold was not included in this analysis because the sample size is too small. Scale bars are all 1 µm.

**Figure 7.**
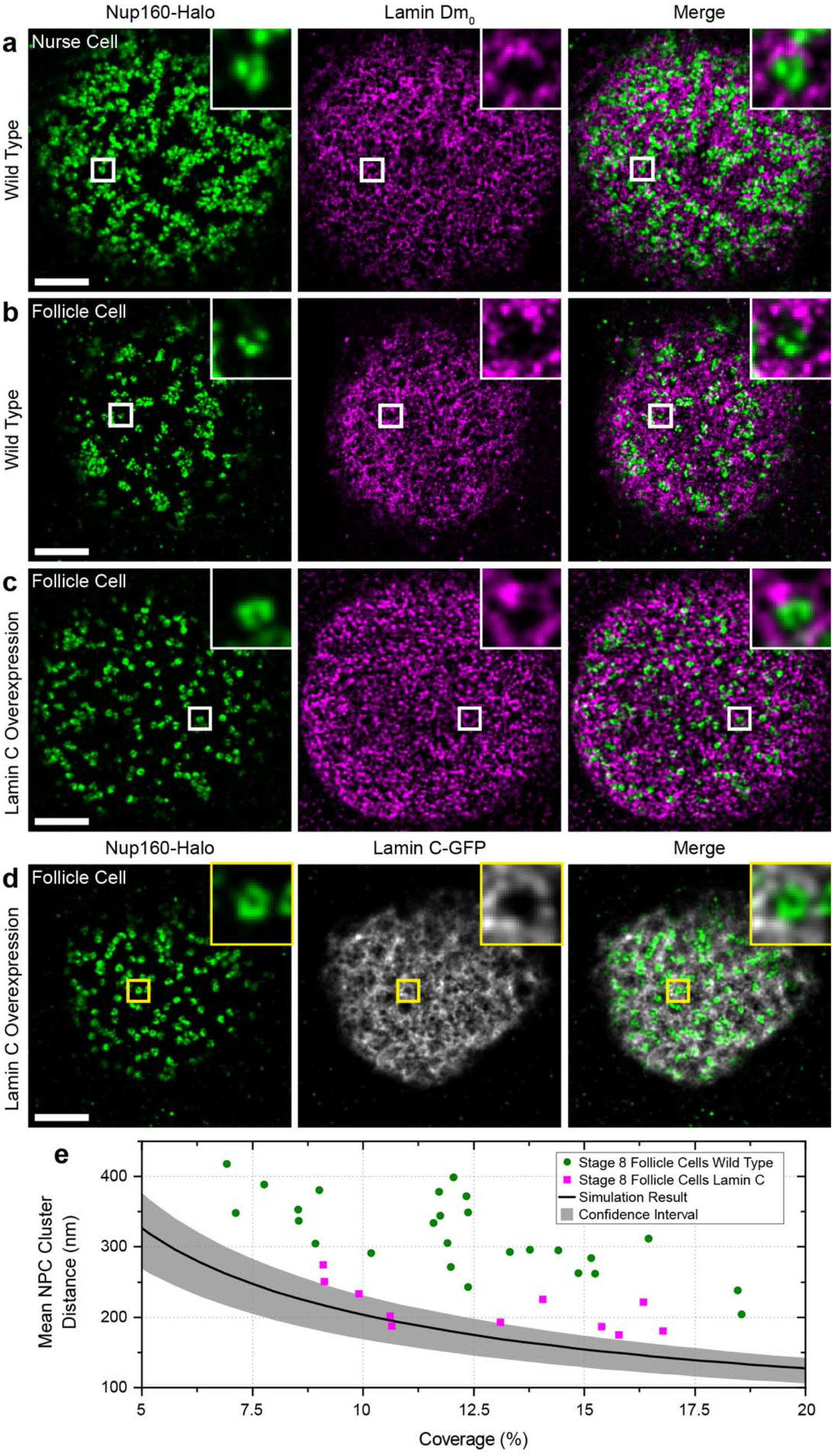
Lamin C inhibits NPC clustering. **a** DNA-PAINT image of Nup160-Halo (green) and Lamin Dm_0_ (magenta) in a wild-type nurse cell. The NPC clusters mainly fall in the gaps in the Lamin matrix. **b** DNA-PAINT image of Nup160-Halo (green) and Lamin Dm_0_ (magenta) in a wild-type follicle cell nucleus. Again, the NPC clusters anti-correlated with Lamin Dm_0_ (column 2). **c** DNA-PAINT image of Nup160-Halo (green) and Lamin Dm_0_ (magenta) in a follicle cell nucleus over-expressing GFP-Lamin C. The NPC are less clustered and Lamin Dm_0_ appears to form shorter and more globular clusters. **d** DNA-PAINT image of Nup160-Halo (green) and GFP-Lamin C (greyscale) in a follicle cell nucleus over-expressing GFP-Lamin C. **e** A graph showing the mean inter NPC cluster distance as a function of the proportion of the nuclear envelope covered by NPCs for wild-type and Lamin C-expressing follicle cell nuclei. The Lamin C expressing nuclei show a more random distribution of NPC. All scale bars are 1 µm.

**Figure 8.**
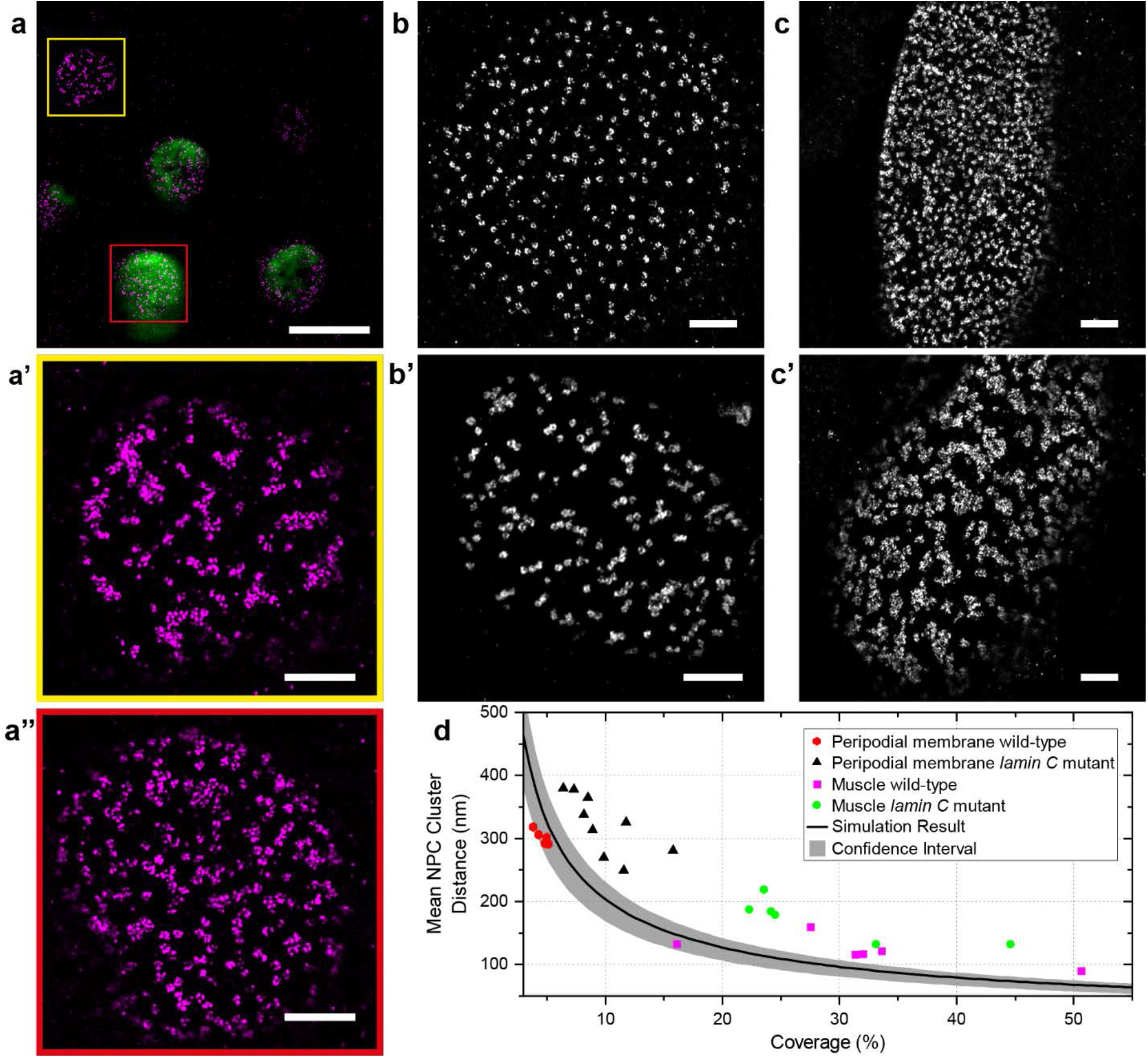
Lamin C Controls NPC Positioning. **a** A DNA-PAINT image of Nup160-Halo (magenta) in a region of the follicular epithelium containing a clone of Lamin C expressing cells (GFP positive, green) adjacent to wildtype (GFP negative) cells. **a’** An enlarged view of the Nup160-Halo distribution in a wild-type nucleus (yellow box in **a**) . The NPCs show normal clustering. **a”** Enlarged views of the Nup160-Halo distribution in a nucleus expressing GFP-Lamin C (detail from red box in **a**). Lamin C overexpression correlates with reduced clustering of the NPCs. **b** A DNA-PAINT image of Nup160-Halo in the nucleus of a wild-type peripodial membrane cell, showing an even distribution of NPC. **b’** A DNA-PAINT image of Nup160-Halo in a nucleus of a peripodial membrane from a *lamin C*^EX5^*/Df(2R) trix* mutant larva, showing clustered NPCs. **c** A DNA-PAINT image of Nup160-Halo in a wild-type larval muscle nucleus, showing an even distribution of NPC. **c’** A DNA-PAINT image of Nup160-Halo in nucleus from a body wall muscle in a *lamin C*^EX5^*/Df(2R) trix* mutant larva, showing clustered NPCs. **d** A graph showing the mean inter NPC cluster distance as a function of the proportion of the nuclear envelope covered by NPCs for wild-type and *lamin C* mutant nuclei a-d. Scale bar in (**a**) is 5 µm. All other scale bars are 1 µm.

### Two colour imaging using DNA- and peptide-PAINT

To test whether one can perform DNA-PAINT and peptide-PAINT simultaneously, we imaged F-actin and Nup160 in stage 3 egg chambers (**Fig. 4a-c**). The Lifeact signal highlights the actin-rich cortex in every cell, revealing the arrangement of the follicle cells around the nurse cells, which lie internally (**Fig. 4c**). Thus, Lifeact-PAINT provides a useful reference system for showing the cell boundaries and locating the imaging plane within the sample, in this case more than 20 µm above the coverslip. We also tested two-colour DNA-PAINT by imaging egg chambers from Nup160-SNAP Gle1-HaloTag homozygous females using two different docking strand oligonucleotides and complementary imager strands labelled with Cy3B and Atto643 (**Fig. 4d-f**). This reveals the localisation of Gle1 in the centre of each NPC surrounded by rings of Nup160 (**Fig. 4g-h**).

### Nuclear pores are clustered in many *Drosophila* tissues

A striking feature of the super-resolved images of nuclear pores (NPCs) in the follicle cells is their organisation into inter-linked chains separated by nuclear pore-free areas, a seemingly non-random distribution. This is surprising, since the nuclear pores are thought to be evenly distributed around the nuclear envelope and previous super-resolution images have revealed an even distribution of NPC in tissue culture cells, which we confirmed in U2OS cells (**Fig. 5j**)^15,19,28-30^. One possibility is that this apparent clustering is a consequence of the fusion of the Halo Tag to Nup160. To test this, we compared the distributions of Nup160-Halo, Nup160-SNAP, Nup188-SNAP and Gle1-Halo in the follicle cells at stage 8 of oogenesis and the nurse cells at stage 4 (**Fig. 5a-h**). Nup160-Halo and Nup160-SNAP show indistinguishable nuclear pore distributions in both the follicle cells and the nurse cells, which have a higher nuclear pore density, but still show a pronounced tendency to cluster (**Fig. 5a-b, e-f**). Furthermore, Nup188-SNAP and Gle1-Halo show very similar distributions of NPCs in both tissues, although they lie nearer the centre of the nuclear pores than Nup160 (**Fig. 5c-d, g-h**). Thus, the non-random distribution of the nuclear pores is not caused by the HaloTag or fusions to Nup160. Finally, we tested whether the clustering is a fixation artefact by imaging Nup160-Halo in living egg chambers after labelling the HaloTag with the cell permeable dye, JF646 ^31^ (**Fig. 5i**). Despite the lower resolution of the Zeiss Airyscan microscope (∼120 nm), the ribbons of clustered nuclear pores are clearly visible, demonstrating that this is the organisation of the nuclear pores in vivo.

To quantify the spatial distribution of nuclear pores we developed an automated pipeline that measures the distances between nuclear pore clusters. As the NPCs become more clustered, the mean distance between clusters increases and this is therefore a proxy measure for clustering. The distances between nuclear pore clusters are also influenced by the density of nuclear pores in the nuclear membrane and their labelling efficiency. To estimate how these two parameters affect the nuclear pore clustering, we performed simulations in which the nuclear pores are distributed randomly without overlapping, but with different densities and labelling efficiencies. Using the data from these simulations, we established a baseline for random nuclear pore distribution along with confidence intervals that can be used as a reference over a wide range of nuclear pore densities and labelling efficiencies (**Supplementary Methods**). To test this analysis pipeline, we imaged Nup96-Halo in U2OS cells, where the nuclear pores are evenly distributed (**Fig. 5j**)^15^. Almost all the measured mean NPC cluster distances in U2OS nuclei fall within the confidence intervals of the simulations, demonstrating that the NPC distribution is close to random. By contrast, the mean inter NPC cluster distances in the follicle cells and nurse cells are significantly larger, indicating that the nuclear pores are more clustered than one would expect by chance (**Fig. 5k**).

To determine whether nuclear pore clustering is a common feature of *Drosophila* cells, we examined the localisation of Nup160-Halo in other tissues. The NPCs in the nuclei of the testis accessory gland and ejaculatory duct of adult males show a similar degree of clustering to the follicle cells (**Fig. 6d, e**). In the 3^rd^ instar imaginal wing disc, the nuclear pores appear clustered in the nuclei of the pouch region (**Fig. 6g**), whereas the peripodial membrane and the fold show no apparent clustering (**Fig. 6h-i**). We also examined the arrangement of nuclear pores in the nuclei of the syncytial blastoderm embryo and observed that most nuclei show some clustering, falling outside the 95% confidence limits for a random distribution (**Fig. 6f, j**). This case also appears in nuclei that have just exited mitosis, indicating that clustering is already present when the nuclear envelope reforms after mitosis (**Supplementary Fig. 9**). Thus, the nuclear pores are clustered at all stages of development and in most tissues analysed, regardless of whether they are dividing progenitor cells or terminally-differentiated post-mitotic cells.

Since the nuclear pores span the nuclear envelope, their organisation could be dictated by either cytoplasmic or nuclear signals. Nuclear pore clustering could be mediated by the cytoskeleton, which interacts with and exerts force on the nuclear envelope both by the direct binding of motors to the nuclear pores and through the LINC complex, which spans the nuclear envelope and can bind to F-actin and microtubules^32-36^. We therefore tested whether either microtubules or F-actin are required for the distribution of nuclear pores by treating egg chambers with the microtubule depolymerising drug, colcemid, or the F-actin depolymerising drug, Latrunculin A. Despite the loss of almost all microtubules or F-actin in treated egg chambers, neither drug had any effect on the distribution of nuclear pores in either the follicle cells or the nurse cells (**Supplementary Fig. 10**).

The nuclear pores are static in the nuclear envelope because they are anchored to the nuclear lamina through multiple interactions between filaments formed by Lamin proteins and specific nucleoporins, making the Lamins good candidates for factors that position the nuclear pores^28,37-39^. The follicle cells and nurse cells express only one Lamin at detectable levels, the B-type Lamin, Lamin Dm_0_^40^. We therefore performed two-colour DNA-PAINT imaging of Lamin Dm_0_ and Nup160-Halo using an anti-Lamin Dm_0_ antibody and a secondary antibody coupled to a docking strand oligonucleotide. This revealed that Lamin Dm_0_ forms a meshwork of filaments along the inner surface of the nuclear envelope of both nurse cells and follicle cells (Fig.7a, b), as reported for other cell types^41,42^. The nuclear pores cluster in the gaps in the meshwork of Lamin Dm_0_ filaments in both cell-types (**Fig. 7a, b**), consistent with the idea that the Lamins organise the nuclear pores.

To test whether the organisation of the nuclear lamina determines the arrangement of the nuclear pores, we mis-expressed the single *Drosophila* A/C-type Lamin, Lamin C, in the follicle cells^43^. This caused the nuclear pores to become more dispersed, with all Lamin C expressing cells having lower mean distances between nuclear pore clusters than wildtype cells with similar nuclear pore densities (**Fig. 7c, d**). Nearly half of the Lamin C expressing cells fall within the 95% confidence limits for random nuclear pore distributions. Lamin C expression also alters the distribution of Lamin Dm_0_, with the Lamin Dm_0_ fibres appearing shorter, thicker and more evenly distributed over the nuclear surface (**Fig. 7c, d**).

To rule out any background variation in nuclear pore distribution caused by the effects of Lamin C expression on egg chamber development or random differences between individual flies, we induced Lamin C expression in mitotic clones in otherwise wildtype Nup160-Halo egg chambers and compared the arrangement of the nuclear pores in the expressing cells versus adjacent non-expressing cells in the same epithelial layer (**Fig. 8a**). The nuclear pores are clustered in the wild-type cells, but are more evenly distributed in the Lamin C expressing cells, confirming that Lamin C acts directly in each nucleus to control nuclear pore positioning (**Fig 8a-a”**).

The observation that Lamin C expression reduces nuclear pore clustering in cells that do not normally express appreciable levels of the protein suggests that *lamin C* mutants should have the opposite effect on Lamin C expressing cells. We therefore examined the organisation of the nuclear pores in the nuclei of the peripodial membrane of the wing disc in *lamin C* ^EX5^/ *Df(2R) trix* larvae, which lack all Lamin C^44^. Although the nuclear pores are evenly distributed in wild-type peripodial membrane nuclei, they are organised into clusters or chains in the *lamin C* null nuclei (**Fig. 8b**). *lamin C* mutants cause muscle defects in both flies and humans, and we therefore also analysed the nuclear pore distribution in nuclei of the larval body wall muscles^45-47^. The nuclear pores are evenly distributed in the nuclear membrane of wild-type muscle nuclei but are highly clustered in the mutant nuclei (**Fig. 8c-d**). Thus, Lamin C is required for the even spacing of the nuclear pores, demonstrating that the composition and organisation of the nuclear lamina controls nuclear pore distribution.

## Discussion

Almost all SMLM imaging to date has been performed on cultured cells that only extend a few micrometres above the coverslip, where one can collect the maximum number of photons with little background signal from out of focus light. Here we describe a simple and cost-effective line scanning microscope system that provides optical sectioning to significantly reduce out of focus background in thicker samples, allowing super-resolution SMLM imaging of tissues. Using this system, we routinely obtain images with a resolution of 30 nm or better at depths of more than 20 µm. This extends the superior resolution of SMLM imaging to a much wider range of specimen and cell-types.

Our microscope works well for various SMLM techniques, including STORM, but we have focused on developing protocols for performing PAINT imaging, because it is immune to photobleaching, allows the use of any fluorophore and is very versatile^48^. For example, we have used peptide-PAINT to obtain high resolution images of the actin cytoskeleton, indirect antibody-mediated DNA-PAINT to visualise the nuclear lamina and direct DNA-PAINT to image endogenous nucleoporins and we demonstrate that these techniques can be combined for two colour, super-resolution imaging. As we generated flies that are homozygous for the tagged proteins, every copy of the protein in the cell carries the tag. This opens up the possibility of extending this approach to count the absolute number of molecules of the protein in any region or structure of interest, using qPAINT ^49^.

One test of a new imaging approach is whether it allows one to detect features that had not previously been visible. Our SMLM system for imaging tissues has revealed that nuclear pores are not evenly distributed in the nuclear envelopes of most *Drosophila* tissues, as they are in tissue culture cells. Uneven distributions of nuclear pores have been previously observed in several contexts. Mutants in some nucleoporins cause clustering of the nuclear pores on one side of the nucleus and a similar phenotype is observed in the absence of Lamin B^50-53^. Secondly, nuclear pore free regions are observed in interphase cells shortly after mitosis and may reflect regions of delayed nuclear envelope reassembly^54^. Finally, there is a specific pathway for inserting new nuclear pores as the nuclear envelope grows during interphase and mutants that disrupt this process lead to zones that are free of nuclear pores as the cell cycle progresses^55-58^. These effects are only observed in specific mutants or cell cycle stages, however, whereas the clustering of nuclear pores in *Drosophila* occurs in wild-type cells, regardless of whether they are still dividing or are postmitotic. More importantly, the non-random distribution of nuclear pores in *Drosophila* occurs on a finer scale than these other phenomena and cannot be detected by confocal microscopy. Since some core nucleoporins have been found to associate with specific chromatin domains, NPC clustering could contribute to the internal organisation of the nucleus, for example, by recruiting specific chromatin regions to the vicinity of the NPC clusters, thereby facilitating the nuclear export of RNAs transcribed from these regions^59,60^.

The localisation of the nuclear pores negatively correlates with the distribution of the Lamin Dm_0_ and is altered by Lamin C expression or loss, indicating that nuclear pore arrangement is determined by the organisation of the nuclear lamina. Biochemical data shows that both B-type and A/C-type Lamins interact with a number of nucleoporins, leading to the suggestion that lamin meshwork anchors the nuclear pores in place^37-39,61,62^. Our results suggest that the B-type Lamin, Lamin Dm_0_, does not directly anchor the nuclear pores, but instead corrals them into the spaces between the Lamin filaments. In support of this view, nuclear pores are clustered in Lamin B free regions in mouse adult fibroblasts (MAFs) that only express Lamin B^62^. Furthermore, in both MAFs and *Drosophila* follicle cells, this clustering is lost when A/C-type Lamins are expressed. Thus, Lamin C seems to play the critical role in determining an even distribution of nuclear pores, either by directly anchoring them to the nuclear lamina, or indirectly, by altering the distribution of the Lamin Dm_0_ filaments.

B-type Lamins are constitutively expressed in both *Drosophila* and mammals, whereas A/C-type Lamins are expressed in a tissue specific manner^43,63^. In mammals, the levels of Lamin A/C rise with increasing tissue stiffness and these form a meshwork in the nuclear lamina distinct from that formed Lamin B filaments that increases the stiffness of the nuclear envelope, thereby protecting it from mechanical deformation^64,65^. It has recently been found that Lamin C expression in *Drosophila* epithelia shows a similar correlation with apical compression^66^. Both the peripodial membrane and the folds of the wing disc are under compression and have high Lamin C levels, whereas the wing pouch epithelium, which is not compressed, has low levels. This is mirrored by the distribution of the nuclear pores in each region, with the peripodial membrane and wing folds having an even distribution of pores, in contrast to the wing pouch, which shows clustered nuclear pores. This strengthens the evidence that Lamin C induces an even distribution of nuclear pores and suggests that this is a response to the mechanical state of the tissue. Since the nuclear pores represent gaps in the nuclear envelope, their redistribution may contribute to the effects of Lamin C on nuclear envelope mechanics, for example if dispersing nuclear pore clusters reduces the risk of rupture under stress. Lamin A/C also affects nuclear mechanics by interacting with lamina-associated chromatin domains (LADs), which increases the stiffness of the nuclear envelope and alters gene expression^67-70^. These effects could also be mediated in part by the altered distribution of nuclear pores through the association of specific chromatin domains with the NPCs^59,60^.

Our observations raise the question of whether the clustered organisation of nuclear pores that we observe in many *Drosophila* tissues is a more general feature of cells in vivo, compared to tissue culture cells. There are very limited data on the distribution of nuclear pores in animal tissues, but it is interesting to note that the nuclear pore arrangement in the follicle cells resembles the ribbons of aligned nuclear pores seen in scanning electron micrographs of Tobacco BY cells ^71^. It will therefore be interesting to determine if this type of nuclear pore organisation is common in other species.

## Methods

### Custom-built slit scanning confocal microscope

A full description of the imaging system employed here may be found in the **Supplementary Methods**. Briefly, a custom line scanning module was attached to the rear port of an inverted microscope base (Olympus, IX83P2ZF) equipped with a 100X 1.35 NA silicone oil immersion objective lens (Olympus, UPLSAPO100XS), motorized xy-sample stage and z-piezo stage. Light propagating from the line scanning module is focused into a line with diffraction limited width and scanned across the field of view with a computer-controlled galvanometer mirror. Fluorescence is collected by the objective lens and relayed onto a camera that has an effective electronic slit which is synchronized with the excitation line being scanning across the field of view. Using a focused excitation line and detection slit provides optical sectioning and reduced out of focus background^72^. Prior to reaching the imaging camera, a combination of dichroic mirrors and emission filters splits the fluorescence emission into two colour channels that are simultaneously imaged on a single camera. The system includes excitation wavelengths at 405, 488, 546, 560 and 642 nm. All aspect of microscope control and data acquisition were carried out using a custom microscopy hardware control platform developed in the LabView environment and freely available on GitHub (**See Code Availability below**). The microscope also includes real-time sample-based drift correction as detailed in the **Supplementary Methods**.

### SMLM Image Reconstruction

Single molecule events on unprocessed camera frames were fitted with a symmetric Gaussian model PSF as described previously^73^. Fitted data were then filtered based on localization precision (0.5-30nm), PSF width (50-150nm), and log-likelihood. The data were then corrected for drift with a redundant cross-correlation algorithm as described previously^74^ and reconstructed to a final super-resolved image.

### Reagents

DNA oligonucleotides (conjugated to fluorophores) were purchased from Integrated DNA Technologies or AtdBio. DNA oligonucleotides conjugated to SNAP or HaloTag ligands were purchased from AtdBio or Biomers.net. A non-commercial modified version of Halo ligand (PBI 300-43) was a gift from Mark McDougall at Promega.

### *Drosophila melanogaster* lines

Single-channel actin images were taken using *w*^1118^ flies. For imaging of nucleopore proteins, several endogenously-tagged lines were created. Nup188, Nup160, and Gle1 were tagged at the C-terminus with SNAP and Halo tags using CRISPR/Cas9 mediated homologous recombination. Guide RNA sequences were identified using http://tools.flycrispr.molbio.wisc.edu/targetFinder/, and cloned into the pCFD3 plasmid for guide RNA expression^75^. Supplementary table 2 lists the guide sites used. Around 1kb of sequence upstream and downstream of the gRNA sites were cloned into the EcoRI and NotI sites of pBluescript SK+ along with either the SNAP or Halo coding sequence, and a short linker sequence, using NEB Hi-Fi assembly mix (New England Biolabs). A mixture of the two guide plasmids (each at 100ng/µl) and homologous repair plasmid (150 ng/µl) was injected into CFD2 embryos ^75^, which express Cas9 under the *nanos* promoter. Resulting F0 adults were screened by PCR to identify potential knock-ins, positives were balanced and stocks created.

UAS-GFP-*lamin C* was a gift from Prof. Kazuhiro Furukawa ^76^. UAS-GFP-*lamin C* was expressed under the control of Gr1-Gal4. Clones of GFP-*lamin C* expressing follicle cells were generated by crossing *y w*; *hs-FLP*; Act5C>CD2>Gal4, UAS:mRFPnls (Bloomington *Drosophila* Stock Center, 30558) to UAS-GFP-*lamin C. lamin C* ^EX5^ (A gift from Prof. Lori Wallrath) and *Df(2R) trix*/*CyO* (BDSC,1896) were both recombined with Nup160-Halo and used to generate *lamin C* mutant third instar larvae.

### Drosophila genetics and dissection of tissues

Standard procedures were used for *Drosophila* maintenance and experiments. Follicle cell clones of Lamin C were induced by incubating hs-FLP; FRT mutant marker adults at 37°C for 2 hours, which were then dissected after 2 days. All flies were fattened on yeast at 25°C for one to two days before dissection. Ovaries were dissected in Schneider’s medium at room temperature and the muscle sheath surrounding the egg chambers was removed. Testis accessory glands and ejaculatory ducts were dissected from adult males, wing discs and body wall muscles were dissected from wandering third instar larvae.

Embryos from 1.5-2 hour collections on apple juice plates were dechorionated with 50% bleach for 3–5 min, and washed three times with water. Embryos were fixed for 20 min in 3% formaldehyde in 0.5X PBS overlaid with heptane (1:1 fixative/heptane). Embryos were transferred to double-sided tape stuck to the base of a 100 mm petri dish, containing 0.2% Tween in PBS. The vitelline membrane was removed using a syringe needle.

### Fixation and Staining

#### Peptide PAINT

Ovarioles were dissected out of the muscle sheath, and warm (38°C) fixation buffer was added (4% methanol-free formaldehyde, 2% Tween 20 in 0.5x Phosphate buffered saline (PBS)). Samples were fixed for 20min at room temperature with rotation. Samples were washed 3x 10min with 0.2% Tween20 in PBS.

Egg chambers were attached to a coverslip (high precision) using Cell-Tak (Corning) and then mounted on concavity slides in approximately 150μl of labelling solution (0.5-5 nM Cy3B Lifeact in PBS with either 20mM sodium sulphite or PCA (3,4 Dihydroxybenzoicmacid), PCD (Protocatechuate 3,4-Dioxygenase Pseudomonas), Trolox ((+-)-6-Hydroxy-2,5,7,8-tetra methylchromane-2-carboxylic acid) – see Supplementary Table 2). Slides were sealed with two-compound silicone glue and kept in the dark for 5-10 min until the glue solidified. Alternatively, egg chambers were attached to the coverslip base of an 8-well chambered coverglass with Cell-Tak, and ∼200-300 µl labelling solution was added. N-terminally labelled Cy3B Lifeact was synthesised by Peptide Protein Research Ltd. Mounting methods, imaging solutions and imaging conditions are indicated in **Supplementary Table 3**.

#### DNA-PAINT

For egg chamber samples with endogenously Halo or SNAP tagged proteins, ovarioles were fixed in 3% formaldehyde in 0.5X PBS for 15 min at room temperature. Samples were washed 3 × 5 minutes in PBS, then permeabilised in 0.5% Triton X-100 for 5 min. They were then quenched with 50 mM Ammonium Chloride in PBS for 5 minutes, washed for 5 minutes in PBS and incubated in Image-iT FX signal enhancer (ThermoFisher Scientific) for 30 min. The labelling reaction was performed for 1 hour at 37°C, with shaking, in 0.5% BSA, 1 µM DTT in PBS with 1 µM of docking oligonucleotide. Samples were washed 3 × 5 min in PBS followed by 0.1% Triton X-100 in PBS overnight. The following day samples were washed 6 × 10 min in PBS before being mounted for imaging. Egg chambers were attached to an 8-well chambered coverglass with Cell-Tak, and ∼200-300 µl imaging solution containing fluorescently labelled imager strand, and oxygen scavengers. Mounting methods, imaging solutions and imaging conditions are indicated in **Supplementary Table 3**.

To combine DNA-PAINT with Lifeact imaging, the protocol was followed as above, but 0.5-2nM Cy3B Lifeact peptide was added to the imager solution.

To perform two colour DNA PAINT imaging of a SNAP or Halo-tagged protein and a GFP-tagged protein, the latter was labelled with a GFP-nanobody coupled to a docking strand oligonucleotide (Massive Tag-q Anti-GFP Kit, Massive Photonics). The same protocol was followed as above, with the GFP-nanobody incubated with the sample along with the SNAP/Halo docking strand.

To perform DNA PAINT with a primary antibody followed by a secondary antibody conjugated to the docking strand, the egg chambers were blocked with 10% bovine serum albumin (in PBS with 0.2% Tween 20) for 1 h at room temperature after fixation and permeabilization. The samples were then incubated with primary antibodies against Lamin Dm0 (1:200) in 0.5% BSA at 4 °C overnight. After 6 × 10 min washes in 0.1 %Tween in PBS, egg chambers were quenched with 50 mM Ammonium Chloride for 5 min. The following steps were the same as the method of single DNA-PAINT described above, except that the secondary antibody conjugated to the docking strand (Massive SDAB 1-Plex, Massive Photonics) was incubated along with the SNAP/Halo docking strand.

For the testis accessory gland, wing discs and body wall muscles, the tissues were fixed with 4% formaldehyde, 2% Tween in 0.5xPBS for 20 min at room temperature. Samples were then washed 3 × 5 min in PBS and then treated with 50 mM Ammonium Chloride in PBS for 5 min. All subsequent steps were the same as for egg chambers, except that the labelling reaction was performed for 2 hours at 37°C, and all steps were undertaken in glass dishes, rather than 1.5ml tubes.

### DNA PAINT on Nup96-Halo U2OS cells

Cells were seeded in a 35 mm imaging dish with a glass bottom 24 h before the labelling, so that they reached 50% confluency next day. Cells were prefixed for 30 s in 2.4% (w/v) PFA in PBS and then permeabilised for 3 min in 0.4% (v/v) Triton X-100 in PBS. Cells were then fixed for 30 min in 2.4% (w/v) PFA in PBS and incubated for 5 min in 100 mM NH_4_Cl in PBS. Cells were then washed 2 × 5 min in PBS and incubated for 30 min in Image-iT FX signal enhancer. The labelling reaction was performed for 2h at room temperature 1µM docking strand with 1µM of DTT in 0.5% BSA in PBS. After the labelling the cells were washed for 3 × 5 min in PBS and imaged in imaging solution.

### Buffers

100x PCD, 40x PCA were made according to Schnitzbauer et al ^77^.

100× Trolox was prepared either according to Schnitzbauer et al (2017), or dissolved directly in Methanol to give a 100 mM solution (as per Cordees et al, 2009^78^, which was diluted to 1 mM in imaging solution and aged using UV exposure to give ∼125 µM TQ and ∼875 µM Trolox before use. This procedure ensured a consistent balance between the oxidizing and reducing capacity of the Trolox. In our hands, 2 minutes of exposure to UV on a gel-doc transilluminator produced the desired ratio.

Basic imaging solution for DNA-PAINT: 1× PBS pH 7.2, 500 mM NaCl.

Basic imaging solution for Peptide PAINT: 1× PBS pH 7.2 500 mM NaCl, or 1× PBS pH 7.2.

### Drug treatments and Immunofluorescence staining

To depolymerise actin and tubulin, egg chambers were treated with 100 μM Latrunculin A and 50 mg/ml Colcemid respectively in Schneider’s medium for 1.5 hours at 25 °C. Control egg chambers were incubated in Schneider’s medium for 1.5 hours at 25 °C. Egg chambers were fixed in 4% Formaldehyde, 2% Tween 20 in PBS for 20 min and then washed 3 × 5 min in 0.1 %Tween in PBS. Egg chambers were then incubated with a 1:500 dilution of Phalloidin-iFluor 405 (Abcam) and anti-α-Tubulin−FITC antibody (Merck) in PBS, 0.5% BSA at 37°C for 1 hour with shaking. Samples were washed 6 x10 min in 0.2% Tween in PBS buffer, mounted in Vectashield with DAPI (Vector Laboratories) and examined via confocal microscopy (Leica SP8).

### Live imaging of Nup160-Halo

Egg chambers were dissected out of the muscle sheath and incubated in Schneider’s medium containing 0.5 µM JF646-Halo for 20 min. The egg chambers were then removed to Schneider’s medium and imaged in an 8-well µ-slide with poly-lysine coating (Ibidi) using a Zeiss Airyscan microscope.

## Data availability

The data that support the findings of this study are available within the article and its Supplementary Information or from the corresponding authors upon request.

## Code availability

Image data was collected with a custom program written in National Instruments (NI) LabVIEW 2016 64-bit, NI DAQmx, and NI Vision Development Module. Image collection software maybe freely accessed on GitHub at https://github.com/Gurdon-Super-Res-Lab/Microscope-Control.

## Acknowledgments

We would like to thank Jonas Ries for providing us with the U2OS cell line expressing Nup96-Halo, Lori Wallrath for sending us the *lamin C* mutant lines, Yongdeng Zhang and Joerg Bewersdorf for the initial idea of building a line scanning microscope and the St Johnston lab for help and advice. This work was supported by a Wellcome Trust Collaborative award (203285), a Wellcome Principal Fellowship to DStJ (080007, 207496), a BBSRC project grant (BB/P026486/1) and by centre grant support from the Wellcome Trust (092096, 203144) and Cancer Research UK (A14492, A24823). E.D. was supported by a Wellcome Trust PhD studentship (109143). J.C. was supported by a Jiangsu Government Scholarship for Overseas Studies (China).

## Author Contributions

G.S. and E.S.A. built the line scanning microscope, designed the data analysis pipeline and analysed all of the image data. J.C. and E.D. developed the DNA-PAINT imaging protocol. E.D. performed the imaging of NPCs in U2OS cells. B.L. developed the Lifeact peptide-PAINT imaging of F-actin. J.C., J.H.R. and E.S.A. performed all other imaging experiments. J.H.R. and A.P. made the endogenously-tagged fly lines. D.St J. acquired funding administered and supervised the project. D.St J., J.C., G.S., E.S.A., J.H.R., wrote the manuscript.

