## Supplementary information for "A method for single molecule localization microscopy of tissues reveals non-random distribution of nuclear pores in *Drosophila*"

#### Supplementary Figures

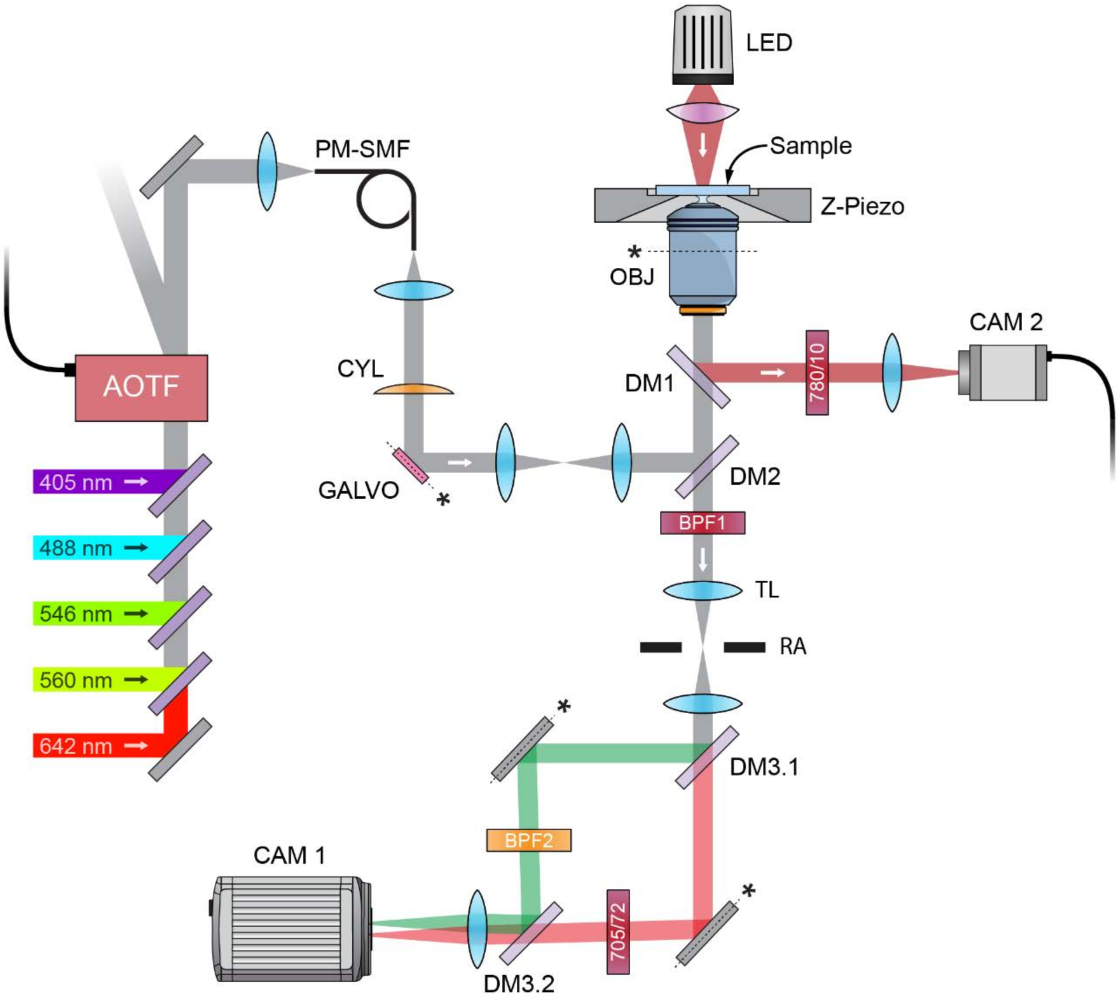


**Supplementary Figure 1. Line Scanning Confocal Microscope Setup.** Five excitation lasers are coupled into a fibre (PM-SMF), collimated at the fibre exit, and pass through a cylindrical lens (CYL) that focuses the beam in one dimension onto a galvanometer mirror (GALVO). The excitation laser light is reflected by a dichroic mirror (DM2) into the objective lens (OBJ) where it forms a line of excitation light at the sample plane. Fluorescence is collected by the same objective lens, separated from the laser excitation light by DM2, passes through a bandpass filter (BPF1) and is imaged by the microscope base's tube lens (TL). A rectangular aperture (RA) is placed at the microscope image plane. The fluorescence image is relayed by a telescope onto the primary imaging camera (CAM1). Fluorescence is split into two colour channels by means of a dichromic mirror (DM3.1), slightly tilted using the subsequent fold mirrors and recombined using a second dichromic mirror (DM3.2). The far-red beam path includes a fixed bandpass filter (705/72) while the alternate beam path includes a motorized filter wheel for easy switching between GFP and Cy3B (BPF2). A far-red lamp (LED) allows for transmitted light imaging using a dichroic mirror (DM1), bandpass filter (780/10) and camera (CAM2) for real-time drift correction during image acquisition. See **Supplementary Methods** for a detailed description of the microscope.

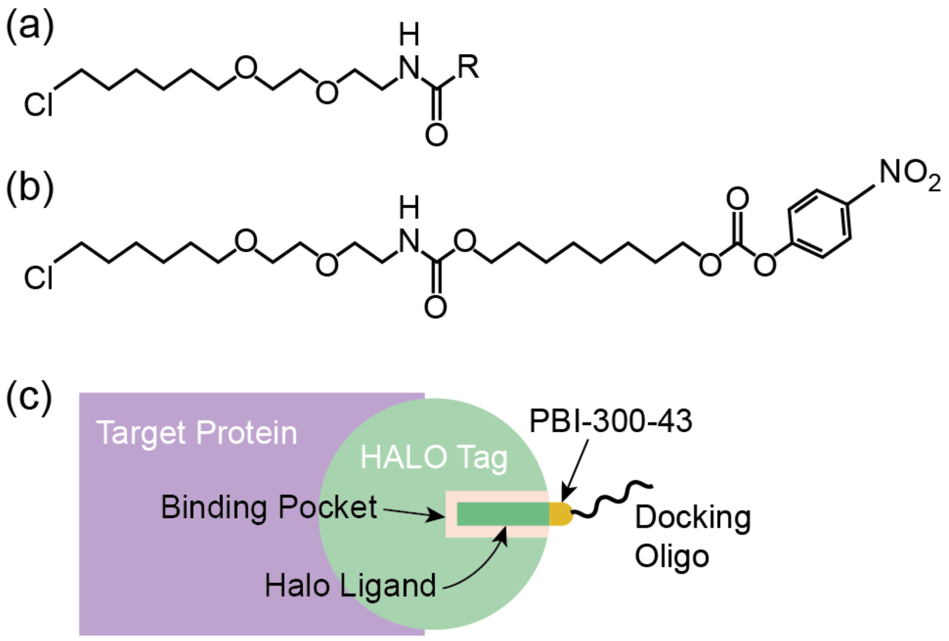


**Supplementary Figure 2**. **Halo ligand schematic. a** Schematic of a typical Halo ligand. **b** Halo ligand modified with PBI-300-43. **c** Geometry of Halo tag and oligo docking strand used for DNA-PAINT.

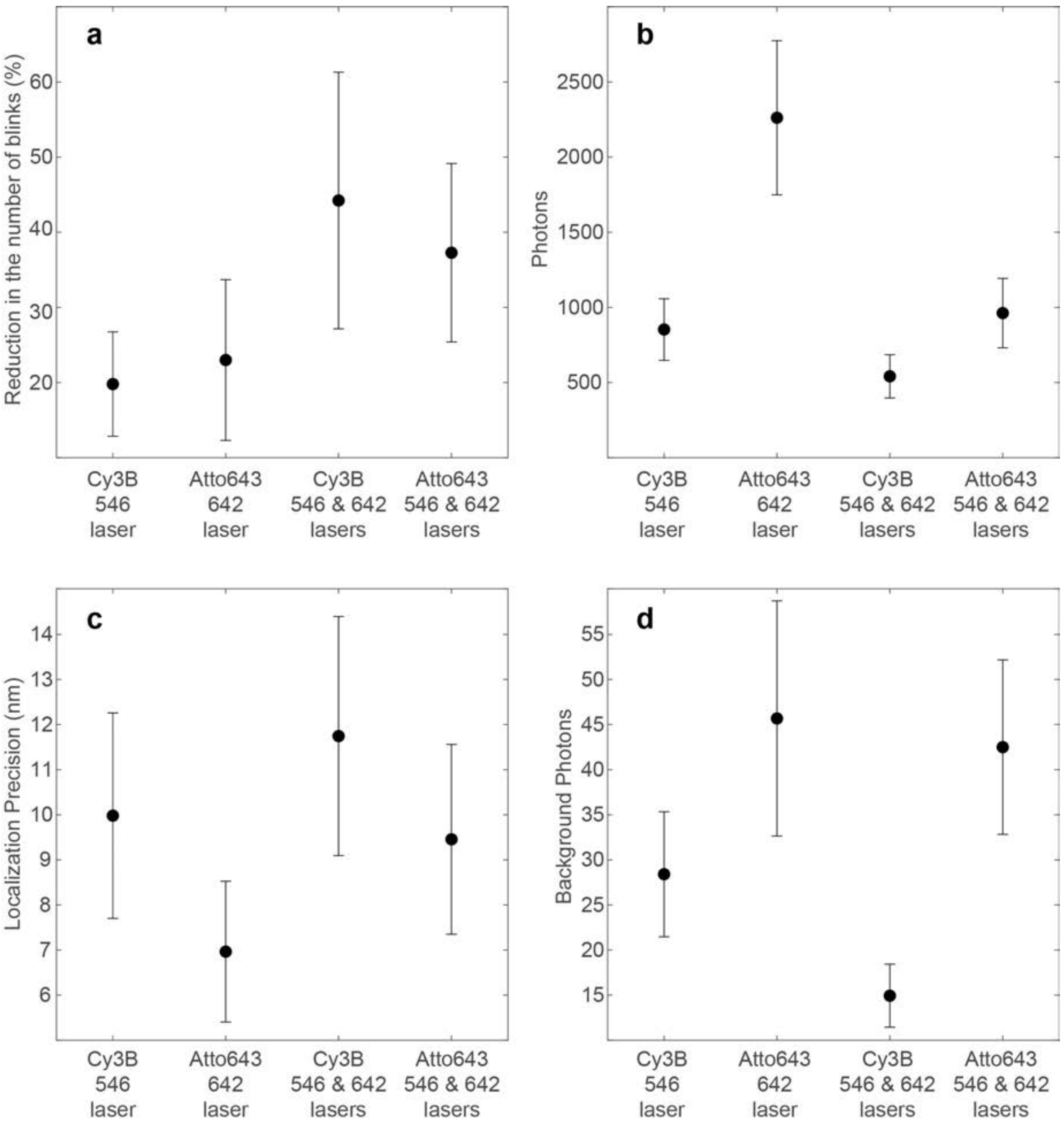


**Supplementary Figure 3: Representative experimental localization imaging results. a** Reduction in the number of blinks calculated between the accumulated number of blinks over a period of 10000 frames at the start of an acquisition and 10000 frames at the end of an acquisition. **b** Mean number of photons**. c** Mean localization precision. **d** Mean number of background photons. X-axis values indicate the fluorophore and excitation lasers used during imaging. Error bars indicate the mean ± the standard deviation (n=6).

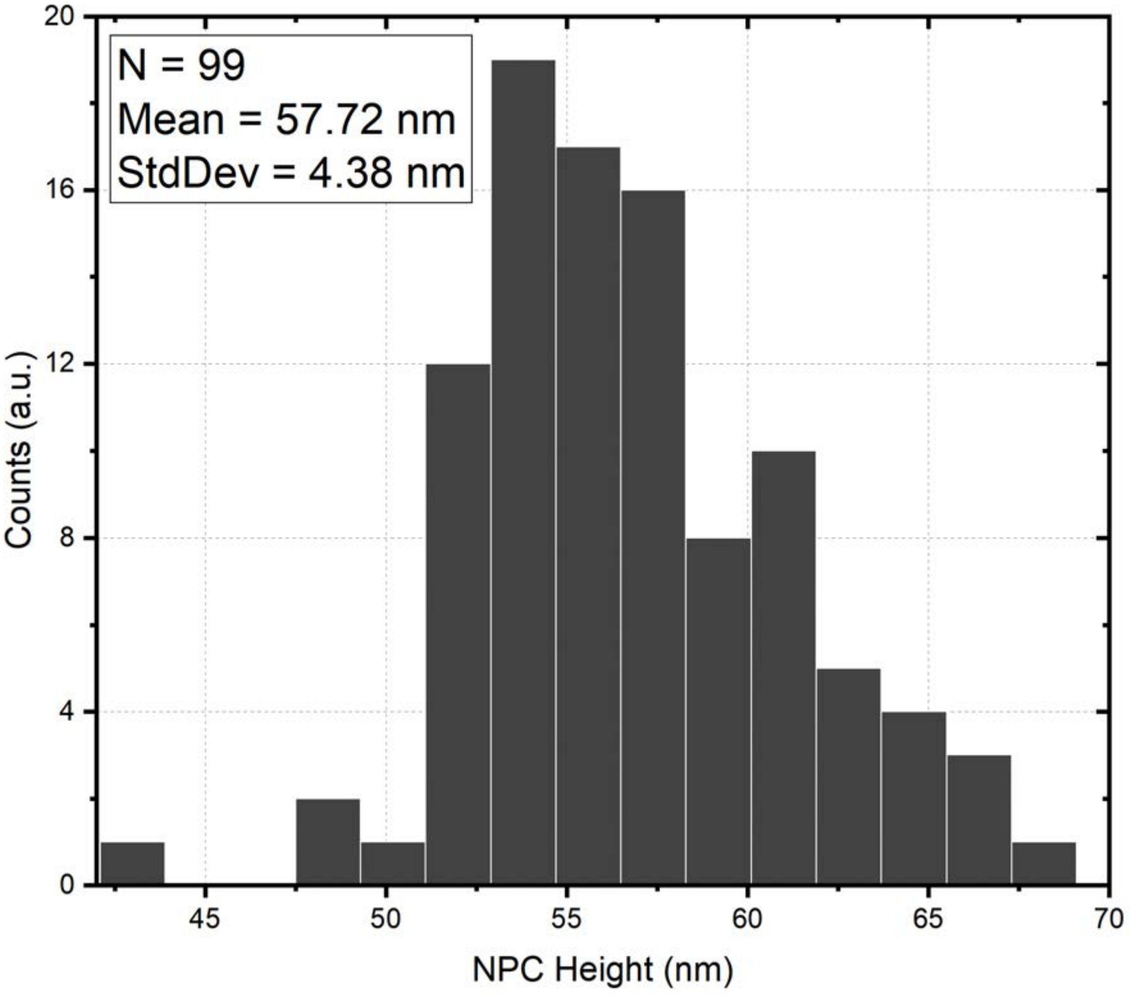


**Supplementary Figure 4. The distribution of measured distances between the nuclear and cytoplasmic rings of the Nup160 NPCs.** 99 ROIs containing cross-sections of NPCs were hand selected from 21 separate Nup160 images. For each ROI, a line was draw between the rings by hand. The ROI and hand annotation were processed by custom scripts in MatLab that rotated the (x, y) localizations such that the long dimension of the NPC rings was parallel with the horizontal axis, and summed along the horizontal axis to generate a histogram with position on the x-axis and number of localizations on the y-axis (example show in main text **Figure 2d**). These data were then fitted with two Gaussians and the distance between the fitted peak positions was determined and plotted on the histogram above.

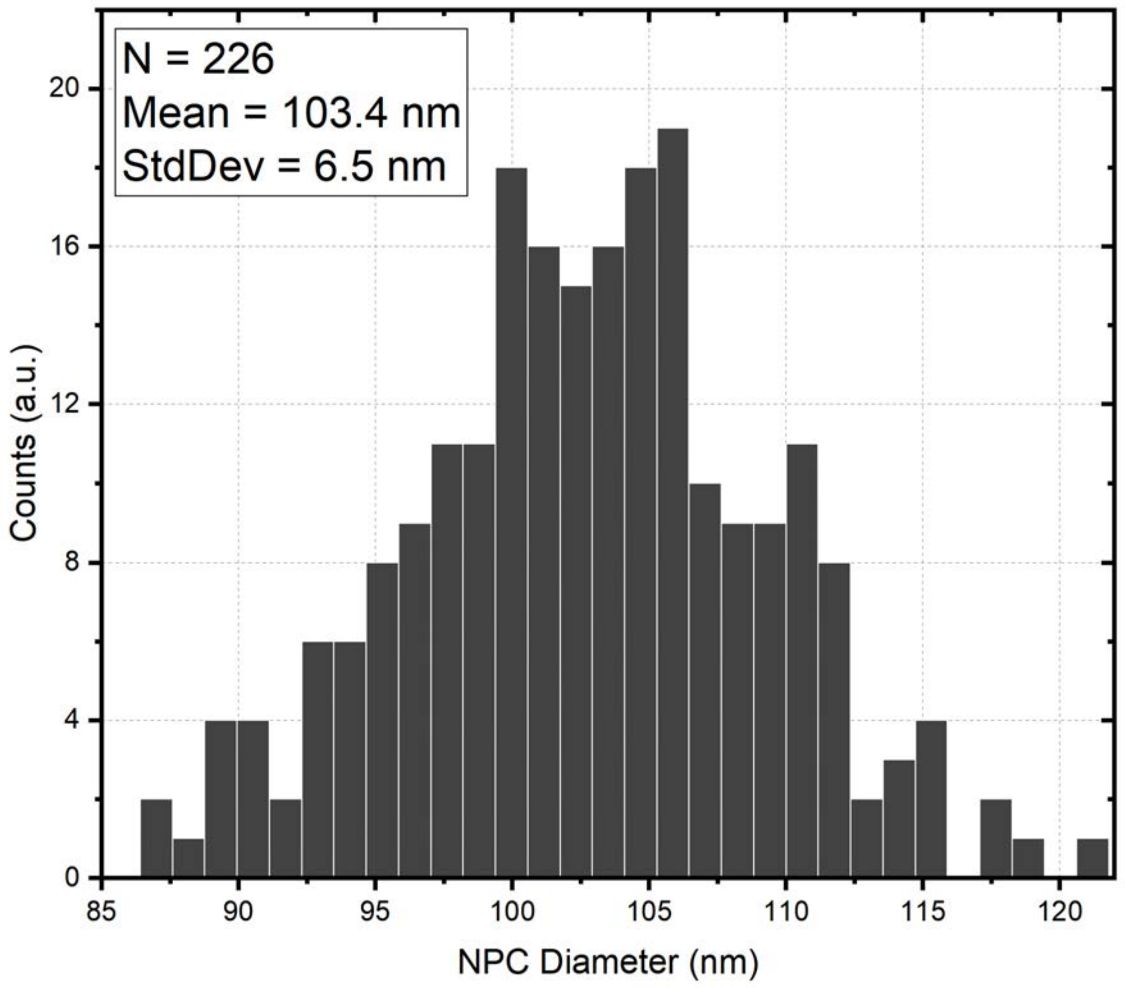


**Supplementary Figure 5. A histogram showing the distribution of measured NPC diameters.** 226 ROIs containing a single, spatially separated, NPC were hand selected from 27 separate Nup160 images. The (x, y) localizations in each ROI were filtered to remove any (x, y) point that was more than σ/8 away from its nearest neighbour. The remaining (x, y) points were fit with a circle to determine the NPC diameter.

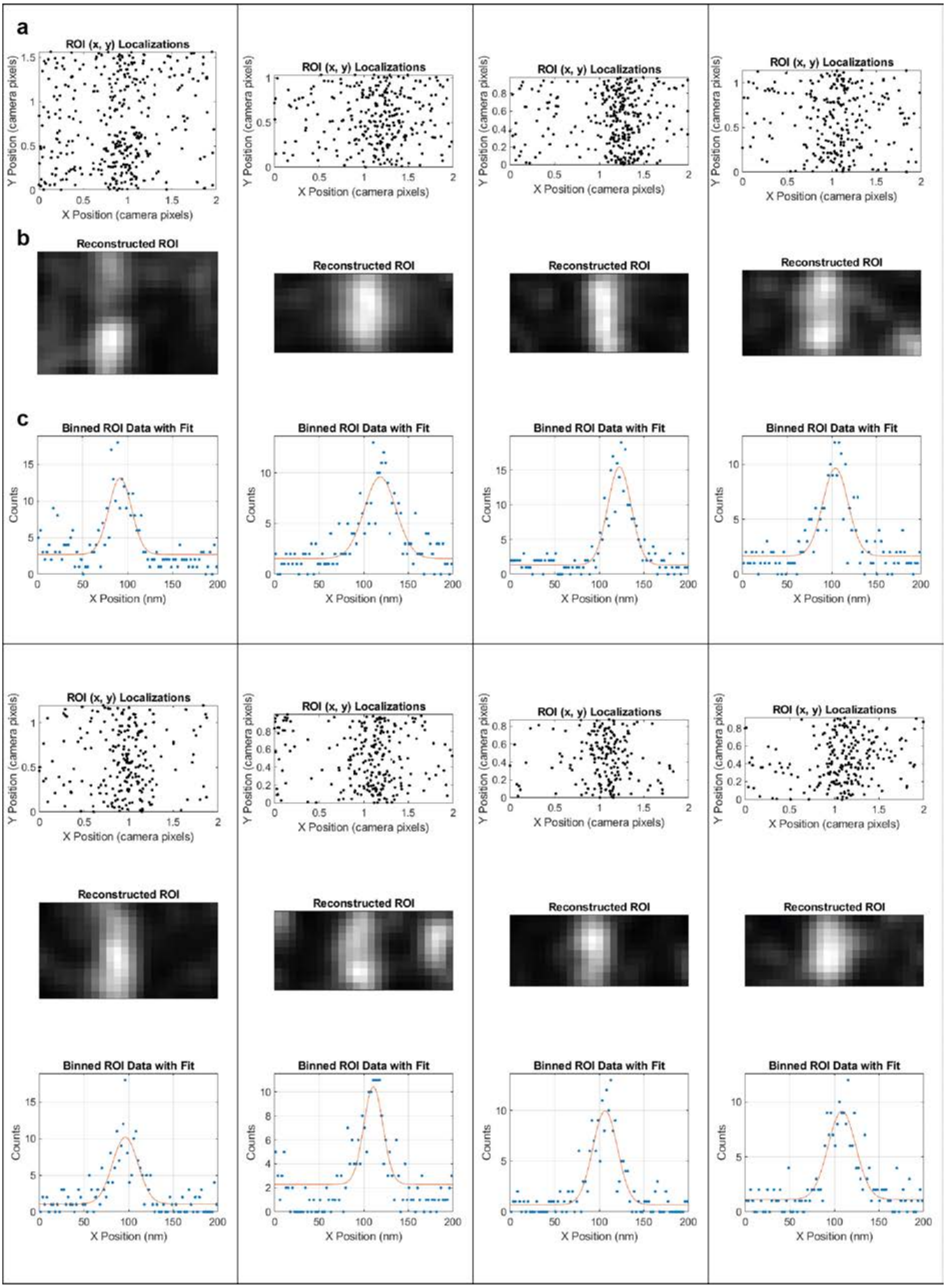


**Supplemental Figure 6. Eight Typical features from actin images binned and fit with a Gaussian. a** Localizations lying along a line, and within a certain distance of the line, are segmented out of the (x, y) localization dataset and rotated such that the filament subset is parallel to the vertical axis. **b** Segmented (x, y) localizations may be reconstructed into a typical SMLM image. **c** Localization are binned (blue data points) and fit (solid red line) with a gaussian to determine the filament FWHM.

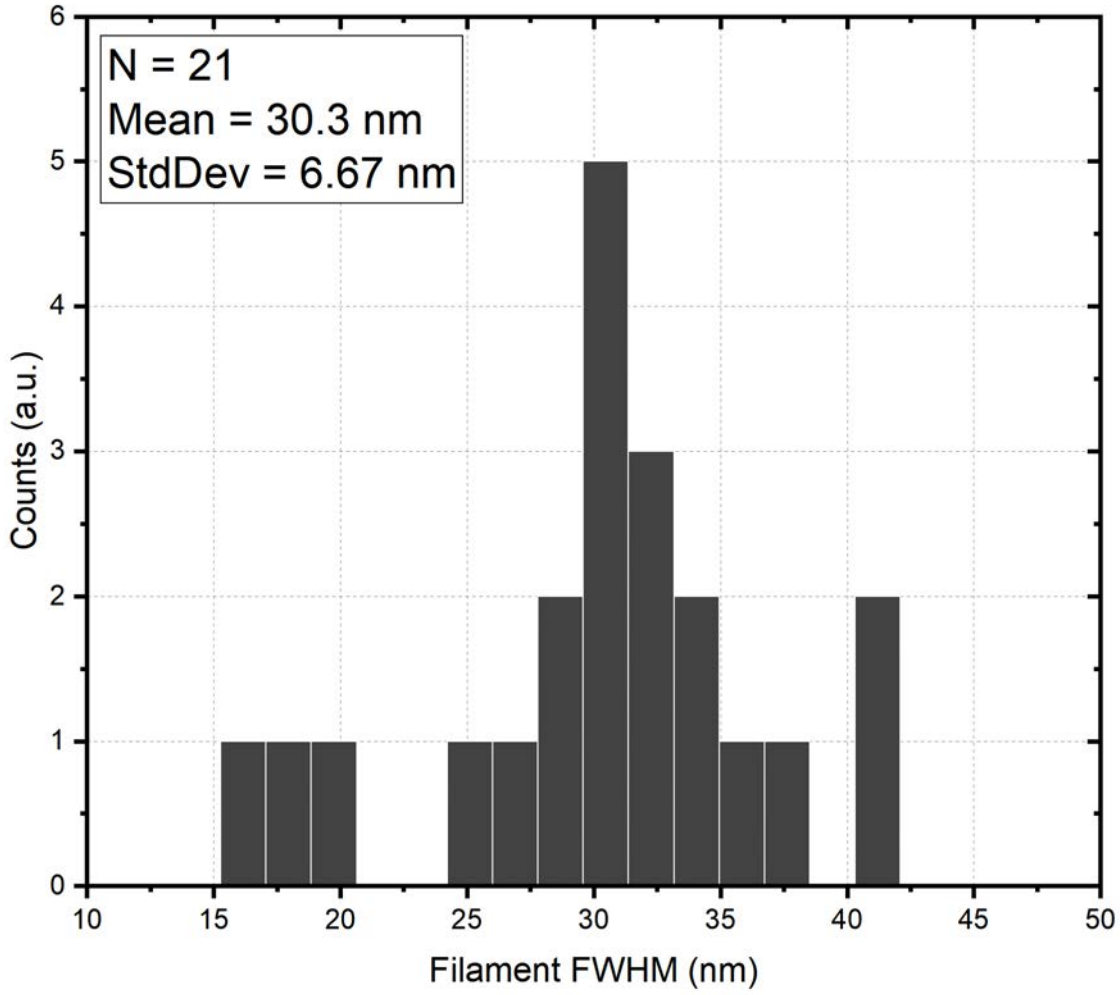


**Supplemental Figure 7. Distribution of fitted FWHM actin feature widths.** 21 actin filaments were hand selected from the three images presented in **Main Text Figure 2** and fit as described elsewhere within the Supplementary Methods. The mean fitted FWHM was 30.3 ± 6.67 (mean ± standard deviation, N = 21) with nearly identical values for features identified in the basal vs. apical images (see **Supplementary Methods** for specific values).

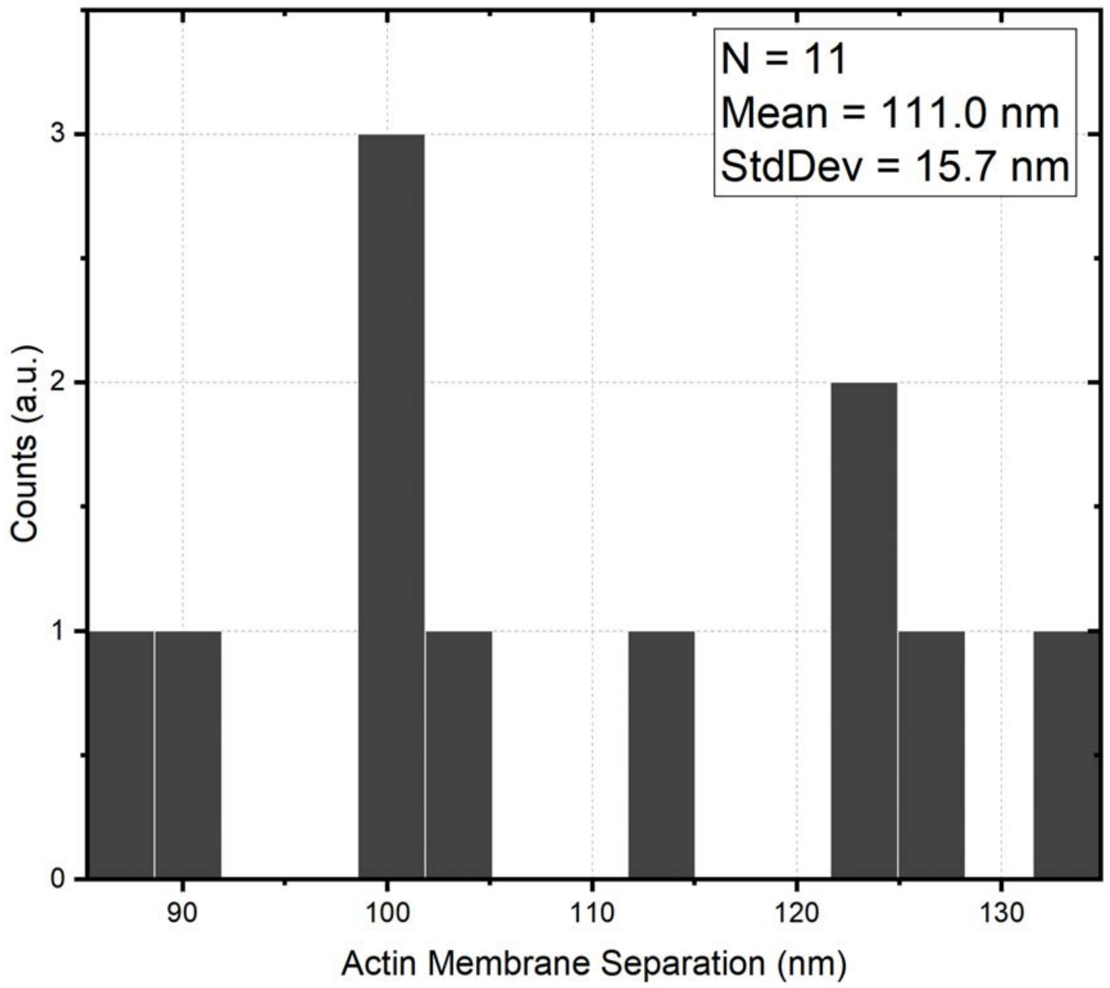


**Supplemental Figure 8. Membrane separation distances as visualized by actin.** 11 locations showing actin cortices in adjacent cells were selected across 7 images. For each location, a line was drawn between the parallel membranes and localizations around the marked lines were segregated, binned, and fit with a double Gaussian function to determine the separation between the two membranes as 111.0 ± 15.7 nm (mean ± standard deviation).

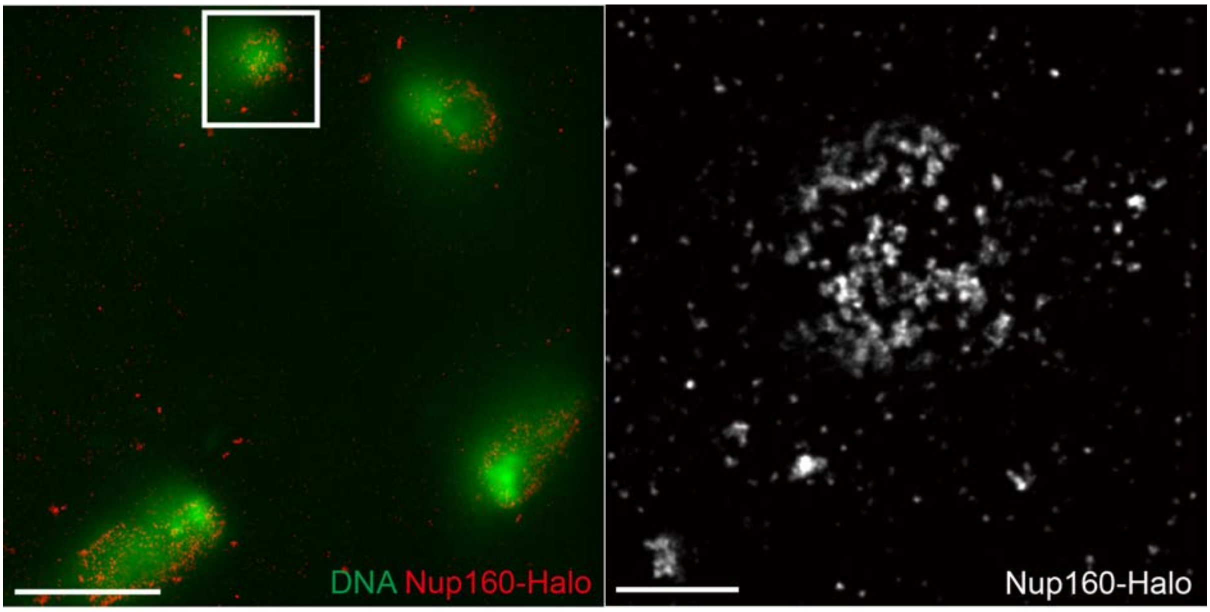


**Supplementary Figure 9. NPC clustering is present in syncytial blastoderm nuclei immediately after mitosis.** Left image, a combined confocal and DNA-PAINT image of DNA (DAPI; green) and Nup160-Halo (red) in a nucleus that has recently divided in a syncytial blastoderm embryo, showing clustered NPCs, scale bar is 5 µm. Right image is a magnified view of the boxed region in (Left) showing NPCs labelled with Nup160-Halo, the scale bar is 1 µm.

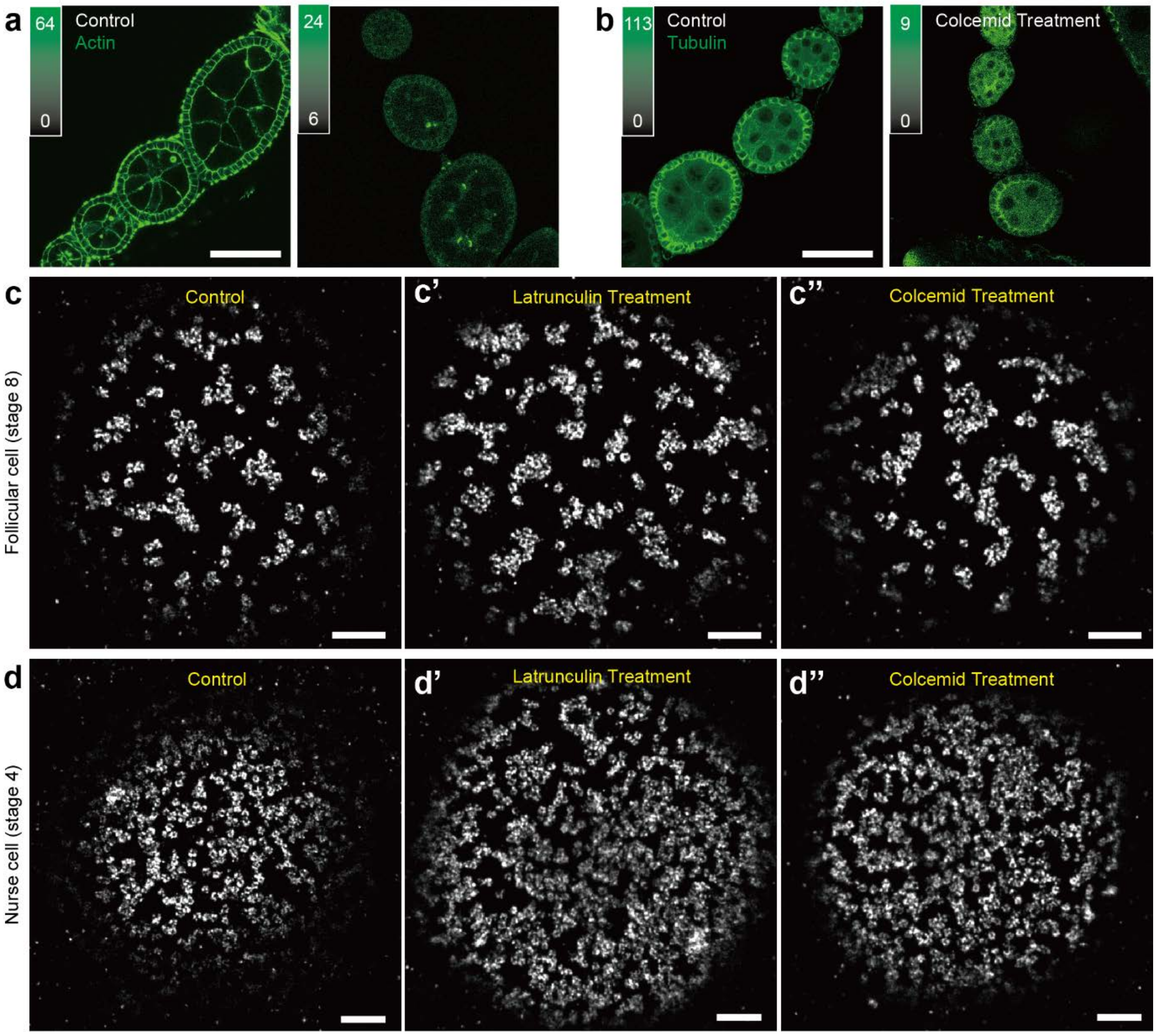


**Supplementary Figure 10. NPC clustering is unaffected by actin or microtubule disruption.** Actin and microtubules were disrupted by treatment with Latrunculin (a) and Colcemid (b), respectively, as shown when imaged with confocal microscopy. In stage 8 *Drosophila* egg chamber, The NPCs (c) in stage 8 follicle cells remain clustered when treated with Latrunculin (c') and Colcemid (c'') and imaged using DNA-PAINT. Similarly, the NPCs (d) remain clustered in stage 4 nurse cells when treated with Latrunculin (d') and Colcemid (d''). Thus, NPC clustering is not dependent on actin or microtubules. Scale bars are 5 µm for panels a and b, and 1 µm for panels c and d.

### Supplementary Tables

#### Supplementary Table 1. Halo tag Effective Labelling Efficiency

| Halo Ligand | Docking Strand | Oligo Storage Time | Fly Genotype | Total No. of NPCs | No. of Gle1 Positive | Percent of Gle1 Positive (%) | Effective Labelling Efficiency (%) |
|---|---|---|---|---|---|---|---|
| **PBI-300-43** | P1 | >1.5 years | Gle1-Halo, heterozygous Nup160-SNAP, homozygous | 210 | 142 | 67.62 | 26.29 |

| | | | | | | | |
|---|---|---|---|---|---|---|---|
| **PBI-300-43** | P1 | >1.5 years | Gle1-Halo, homozygous Nup160-SNAP, homozygous | 724 | 611 | 84.39 | 20.72 |
| **O2** | P3 | < 1 month | Gle1-Halo, heterozygous Nup160-SNAP, homozygous | 194 | 126 | 64.95 | 24.56 |
| **O2** | P1 | < 1 month | Gle1-Halo, heterozygous Nup160-SNAP, homozygous | 674 | 578 | 85.76 | 43.24 |

#### Supplementary Table 2. Guide Sites

| | Recognition sequence |
|---|---|
| **Nup160** | |
| Guide1 | ggccgttctgcagatgcgaa**cgg** |
| Guide2 | gcagatgcgaacggatcttcagg |
| **Gle1** | |
| Guide 1 | gtgcaatctgccgctcacgc**agg** |
| Guide 2 | gccacaactttattgattga**agg** |
| **Nup188** | |
| Guide 1 | ttcggttgtaaagtaattca**tgg** |
| Guide 2 | cgactgagctgctagccctt**cgg** |

#### Supplementary Table 3. Imaging conditions

| **Figure Number** | **Imager Stand and Concentration** | **Docking strand(s)** | **Slide Type** | **Oxygen Scavengers** | **Frame Rate (FPS)** | **No. of Frames** | **Laser Power (just before the Objective)** |
|---|---|---|---|---|---|---|---|
| Fig. 2 | P3-Cy3B 0.5 nM | Halo-P3 | 8-well chambered coverglass | PCA PCD Trolox | 4 | 60,000 | 1.1 mW (546 Laser) |
| Fig. 3a | Cy3B Lifeact 5 nM | -- | Concavity slide | 20 mM Sodium Sulfite | 8 | 100,000 | 1.52 mW (546 Laser) |
| Fig. 3b | Cy3B Lifeact 1 nM | -- | Concavity slide | 20 mM Sodium Sulfite | 8 | 100,000 | 1.52 mW (546 Laser) |
| Fig. 3c | Cy3B Lifeact 1 nM | -- | 8-well chambered coverglass | PCA PCD Trolox | 8 | 60,000 | 0.87 mW (546 Laser) |
| Fig. 4a-c | Cy3B Lifeact 1nM and P1-Atto643 1nM | Halo*-P1 | 8-well chambered coverglass | PCA PCD Trolox | 4 | 60,000 | 0.87 mW (546 Laser) and 2.47 mW (642 Laser) |
| Fig.4d-h | P2-Cy3B 1nM and P1 Atto643 0.5nM | Halo*-P1 and SNAP-P2 | 8-well chambered coverglass | PCA PCD Trolox | 4 | 60,000 | 0.87 mW (546 Laser) and 2.47 mW (642 Laser) |
| Fig. 5a-h | P1-Atto643 0.2nM | Halo*-P1 | 8-well chambered coverglass | PCA PCD Trolox | 4 | 60,000 | 2.47 mW (642 Laser) |
| Fig. 5j | P1-Cy3B 2 nM | Halo*-P1 | 35 mm glass-bottomed dish | Oxidase, Catalase, Glucose | | 25,000 | Imaged on Nikon N-Storm system. |
| Fig. 6 | P1-Atto643 0.5nM | Halo*-P1 | 8-well chambered coverglass | PCA PCD Trolox | 4 | 60,000 | 2.47 mW (642 Laser) |

| | | | | | | | |
|---|---|---|---|---|---|---|---|
| Fig. 7 | P1-Atto643 0.2nM | Halo*-P1 | 8-well chambered coverglass | PCA PCD Trolox | 4 | 60,000 | 2.47 mW (642 Laser) |
| Fig. 8a | P1-Atto643 0.2 nM | Halo*-P1 | 8-well chambered coverglass | PCA PCD Trolox | 4 | 60,000 | 2.47 mW (642 Laser) |
| Fig. 8b-c | P1-Cy3B 0.8 nM | Halo-P1 | 8-well chambered coverglass | PCA PCD Trolox | 4 | 60,000 | 0.86 mW (546 Laser) |
| Supplementary Fig. 4 | P1- Atto643 0.2 nM | Halo*-P1 | 8-well chambered coverglass | PCA PCD Trolox | 4 | 60,000 | 2.47 mW (642 Laser) |
| Supplementary Fig. 5c-d | P1 -Atto643 0.5 nM | Halo*-P1 | 8-well chambered coverglass | PCA PCD Trolox | 4 | 60,000 | 1.1 mW (642 Laser) |

Note: * represents that the docking strand is conjugated with PBI-300-43 halo ligand. fps: frame per second.

### Supplementary Methods

#### Custom-built slit scanning confocal microscope

A shown in **Supplementary Figure 1**, five CW excitation lasers at 405 (Coherent, Obis 405nm LX), 488 (Coherent, Obis 488nm LS), 546 (MPB Communications, 2RU-VFL-P-1000-546-B1R), 560 (MPB Communications, 2RU-VFL-P-2000-560-B1R) and 642 nm (MPB Communications, 2RU-VFL-P-2000-642-B1R) are combined with respective dichroic mirrors and pass through an AOTF (AA Opto-Electronics, AOTFnC-400.650-TN) allowing excitation power control. The diffracted beam or beams from the AOTF are coupled into a polarization-maintaining single mode fibre (PM-SMF; Thorlabs, PM-S405-XP-Custom). Upon exiting the fibre, the diverging light is collimated using a 4X objective lens (Olympus, Plan N 0.10NA; available from Thorlabs as RMS4X) and passes through an adjustable rectangular aperture (Owis, 27.140.0707). Next, the collimated beam passes through a cylindrical lens (CL; Edmund Optics, 68-161). The cylindrical lens focuses the collimated light, along one axis, onto a galvanometer (galvo) mirror (GALVO; ScanLabs, dynAxis XS 7-1) that is conjugated to the pupil plane of a 100X 1.35 NA silicone immersion objective lens (OBJ; Olympus, UPLSAPO100XS) mounted in an inverted microscope base (Olympus, IX83P2ZF). Excitation light reflecting from the galvo mirror enters the microscope base from the rear port, lower deck, and is reflected to the objective lens using a dichroic mirror (DM2; Chroma, 59007bs or ZT405/488/561/647rpc) mounted in the motorized filter turret of the microscope base. The resulting excitation light is focused along one dimension and collimated along the other, when it reaches the objective back pupil plane. The extent (width or diameter) of the light entering the objective back pupil plane is chosen to ensure the pupil is overfilled by the collimated dimension. In this arrangement, the objective lens focuses the excitation light into a line at the sample plane with a diffraction-limited width. The height of the excitation line is set by the rectangular aperture mentioned above prior to the cylindrical lens. In this way, a line with diffraction-limited width and height of approximately 20 µm is created at the sample. This line of excitation light may be scanned along one dimension, orthogonal to the long axis of the excitation line, by movement of the galvo mirror. Fluorescence emission from the sample is collected by the same objective lens and separated from the laser excitation light with a dichroic mirror (DM2). After the dichroic mirror, fluorescence emission passes through a multi-channel bandpass filter (BPF1, Chroma, 59007m or ZET405/488/561/647m) inside the microscope body. Emission light leaves the microscope base via the side port. An adjustable rectangular aperture (RA;

Owis, 27.140.0707) is placed outside the microscope base at the image plane formed by the tube lens. This aperture limits light in the detection path to fluorescence emission from the target field of view in the sample. Next, fluorescence emission is split into two colour channels with a dichroic mirror (DM3.1; Chroma, ZT647rdc). The separated emission light follows identical paths. Each path includes a mirror conjugated to the objective back pupil plane allowing the emission to be slightly offset without losing telecentricity on the detector. The far-red path includes a bandpass filter (Chroma, ET705/72m). The other path includes a motorized filter wheel with two bandpass filters (BPF2). The first (Chroma, ET525/50m) may be used for DAPI/GFP type detection while the second (Chroma, ET590/50m) may be used for Cy3B/ATTO565 type detection. The separated emission paths are recombined with a dichroic mirror (DM3.2) and imaged onto an sCMOS camera (Hamamatsu, Orca-Flash 4.0 V3). The effective pixel size on the camera is 98 nm. A slight tilt is added to the respective mirrors in the separated path such that two images are arranged next to each other on the camera chip, one for the fixed far red channel and the other for the orange or green channel. The sCMOS camera is operated in so-called "light sheet mode". This mode takes advantage of the inherent "rolling shutter" on an sCMOS camera to reduce the number of exposed pixels to one or more rows at any given time. In this mode, a row of pixels is exposed for a predefined exposure time. When that exposure has finished the next row of pixels is exposed for the same time. This process is repeated such that subsequent rows of pixels are sequentially exposed across a predefined region of interest (ROI) on the camera chip. The per-row exposure time may be user set and, independently, how quickly new rows are exposed may also be user set. In this way, the number of adjacent rows simultaneously being exposed may be controlled by the user. Limiting the camera exposure to a set number of rows is equivalent to using a mechanical (confocal) slit to limit out-of-focus background from reaching the detector[1]. As the exposed rows sequentially progress along the camera chip, the glavo mirror synchronously scans the excitation line across the sample. In this way, background signal is greatly reduced when imaging in thick specimen away from the cover glass. Synchronization between the galvo mirror and sCMOS camera is carried out with a data acquisition card (National Instruments, PCIe-6343) and custom software written in the LabView environment (National Instruments, LabView 2016). All aspects of microscope control and data acquisition are carried out using a custom microscopy hardware control platform developed in the LabView environment and freely available on GitHub (https://github.com/Gurdon-Super-Res-Lab/Microscope-Control).

#### Real-time axial (z) sample drift correction

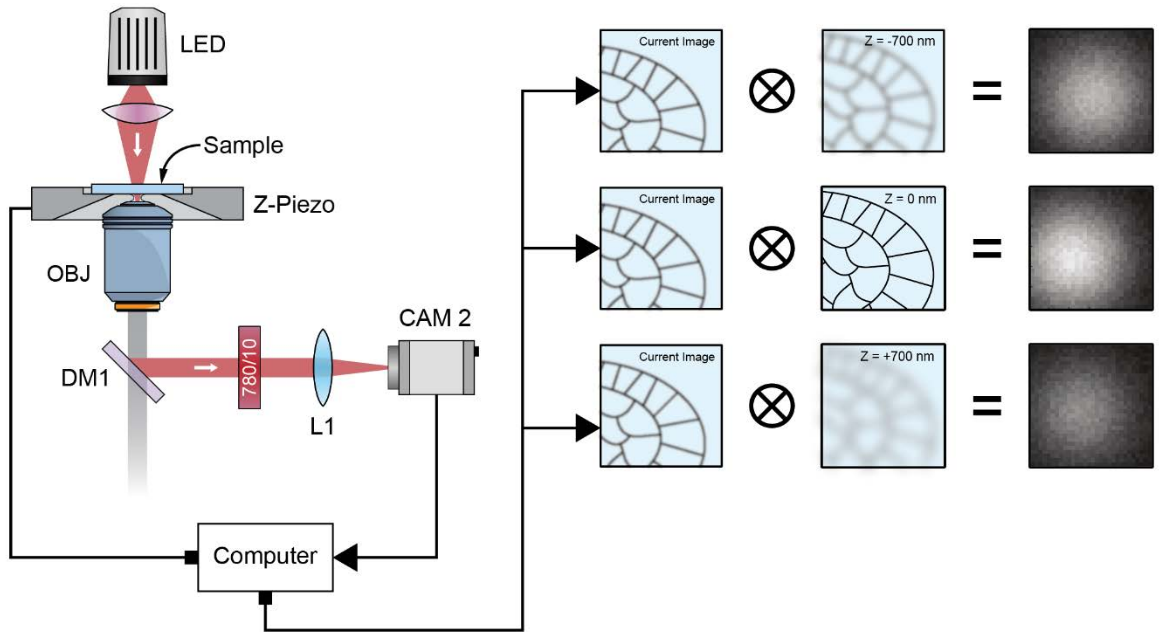


**Supplementary Figure 11. Real-time drift correction setup.** Far-red 850 nm light is emitted from an LED lamp (LED) mounted above the sample. The LED lamp light is collected by the objective lens (OBJ) and separated from the fluorescence beam path with a dichroic mirror (DM1). Light reflected from DM1 passes through a bandpass filter (780/10) and is focused by a 100 mm focal length lens (L1) onto a USB camera (CAM2). The real-time, or current image, collected by CAM2 is cross correlated with previously collected reference images at, above and below the focal plane to track the (x, y, z) sample position. The computer uses the cross-correlation information to correct for any sample Z drift by adjusting the position of the sample pizeo z-stage (Z-Piezo) during image acquisition.

Unlike cultured cells, thick specimens such as *Drosophila* egg chambers may not tightly adhere to a slide or cover glass surface and the desired focal plane is typically many micrometres above the cover glass. These two complications preclude using two of the most common and robust sample drift correction methods: (1) tracking the axial cover glass position with a reflected far red beam and (2) attaching fiducial markers to the cover slip for lateral drift correction in post processing. The most critical of the two is the requirement that the sample z-position stay constant throughout image acquisition. Axial sample drift of just a few hundred nanometres will cause postprocessing lateral drift correction methods to fail. In fact, in early experiments, axial sample drift on the order of tens of micrometres was regularly observed during imaging acquisition (3 to 4 hours) rendering the final image unintelligible. Thus, following from the work of McGorty and co-workers[2], we implemented a **real-time** drift correction scheme based on measurements of the sample itself, not the coverslip or fiducial markers. For this reason, as shown in Supplementary **Figure 11**, the microscope setup was modified to include a light emitting diode (LED) lamp (Thorlabs, M850LP1) centred at 850 nm for transmitted light imaging of the sample outside the normal fluorescence emission range. An additional short-pass dichroic mirror (DM1; Chroma, ZT775sp-2p) was placed just below the microscope objective lens to reflect this light out of the system, through an emission filter (Thorlabs, FL780-10) and tube lens (Thorlabs, AC254-100-B), and onto an inexpensive sCMOS camera (Edmund Optics, 84-933). In this way, transmitted light images at approximately 780 nm could be collect while simultaneously collecting fluorescence images. Prior to starting fluorescence image acquisition, a short z-stack of 11 transmitted light images spaced 100 nm apart in Z, centred around the initial sample position, were collected using the lamp and camera as described above. The central and two extreme images were marked as reference images and used to calculate a dimensionless parameter $\xi$ following from McGorty and co-workers[2] as $\xi_n = (PV\{C_{+,n}\} - PV\{C_{-,n}\})/PV\{C_{0,n}\}$ where $PV\{C_{+,n}\}$ is the peak correlation value between the current image and the upper most reference image, $PV\{C_{-,n}\}$ is the peak correlation value between the current image and the bottom most reference image and $PV\{C_{0,n}\}$ is the peak correlation value between the current image and the central reference image. The stack of reference images was used to generate a z-position vs. $\xi$ curve which was fit with a linear function and used as a calibration. After the transmitted light reference images were collected and the calibration curve was generated the drift correction started automatically. Specifically, a transmitted light image would be collected approximately 1 time per second (approximately 40 msec exposure time and 20 frame accumulations) and used to compute $\xi$ for that image. If the sample Z position was found to have changed by more than ±20 nm, the drift amount would be fed back to the sample z-piezo stage. The sample z-piezo stage would then be stepped by the drift amount, in the opposite direction, to return the sample to its original axial position. In this way, the desired focal plane was maintained by directly imaging the sample. Although this method allows the position of the sample to be tracked in XY and Z, stepper motor driven sample XY stages do not allow small enough repeatable steps to make XY correction reliable while imaging. We therefore adopted a hybrid approach. While imaging, the sample Z position was actively tracked and corrected 1 to 2 times per second using the piezo sample Z stage (Märzhäuser, 00-55-550-0800). SMS (super resolution) camera frames were collected in blocks (cycles) of 1000. After each cycle (1000 frames) the collection of camera frames would pause and if the sample was found to have drifted more than a user pre-set amount, the sample XY stage would be appropriately stepped until the sample's original XY position was restored. In this way, during a 3 to 5 hour image acquisition session the sample's XY position would be maintained adequately enough for the sample's Z drift correction to continue functioning and would not cause any unexpected jumps in sample position while recording camera frames that would otherwise cause the post-processing XY sample drift correction method to fail. The LabView microscope control package detailed above was used to collect the transmitted light images, generate the sample z-calibration curve and fit, and perform the closed loop z-piezo stage control.

##### Postprocessing lateral (x, y) drift correction

For postprocessing lateral (x, y) drift correction, the algorithm from Wang and co-workers[3] was used. The MatLab drift correction function available on the authors GitHub page (https://github.com/yinawang28/RCC) was used with slight modifications to work with the fitting data from our analysis pipeline.

##### Two colour image registration

Two colour images were registered using the method previously described by Huang and co-workers[4]. In brief, Tetraspeck beads immobilized on glass and mounted in index matched media were raster scanned across the microscope field of view. At each step in the scan, an image was recorded in both colour channels. Beads positions in each recorded frame were localized and the positions saved. Corresponding bead positions in each colour channel were used to generate a transformation matrix that was applied to the super-resolution images. Typical registration error between bead images was less than 5 nm. Image registration calibration was performed daily.

#### Morphological Image Analysis

###### Image binarization

Circular regions of interests from grayscale super-resolved images were binarized using an adaptive threshold based on the gaussian weighted mean intensity of an 11-pixel neighbourhood around each pixel. For images labelled for Nup160, isolated and spur pixels were removed from the resulting binary image along with foreground areas with sizes smaller than half the expected area of a Nuclear Pore Complex (NPC). Next, the binary image was morphologically closed with a square structuring element of 2 pixel, followed by an opening operation with the same structuring element. Finally, areas of the binary image where the mean intensity in the corresponding grayscale image was less than 10% of the maximum grayscale value were considered as nonspecific signal and removed. Results for a variety of coverage scenarios ranging from low to high coverage are illustrated in **Supplementary Figure 12.**

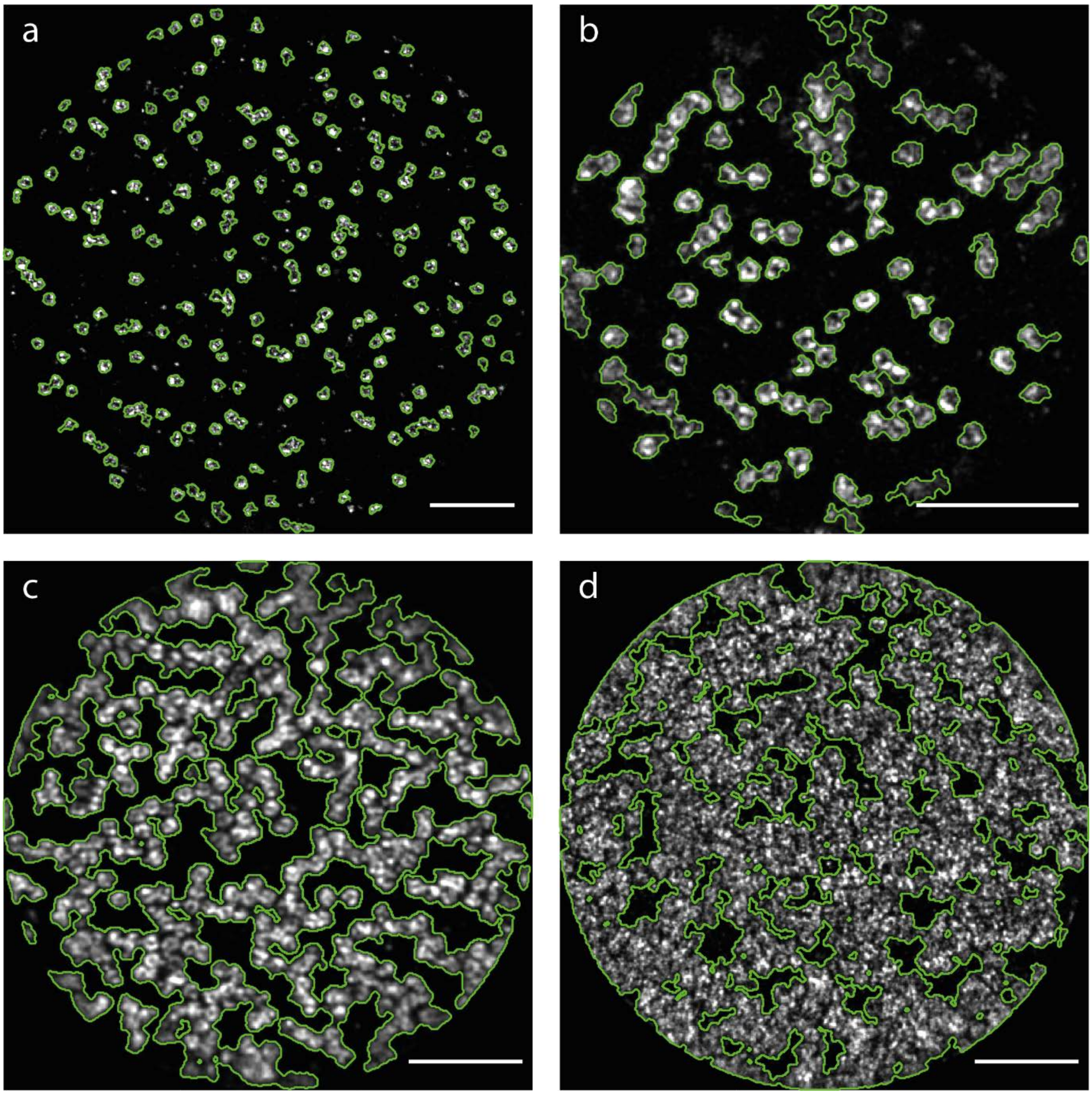


**Supplementary Figure 12: Binarization of super-resolved images**: Green lines indicate the boundaries between black and white pixels in binarized super-resolved images and are super-imposed on the original greyscale image for comparison purposes. Scale bars: 1 µm (**a**) A binarized super-resolved image of NPCs in the peripodial membrane of the *Drosophila* wing disc. The NPCs cover 5% of the nuclear membrane. (**b**) A binarized super-resolved image of NPCs in a follicle cell in a *Drosophila* stage 8 egg chamber ectopically expressing Lamin C. The NPCs cover 13% of the nuclear membrane. (**c**) A binarized super-resolved image of NPCs in a nurse cell in a *Drosophila* stage 3 egg chamber with 55% NPC coverage of the nuclear membrane. (**d**) A binarized super-resolved image of NPCs in a nurse cell in a *Drosophila* stage 5 egg chamber with 75% NPC coverage of the nuclear membrane.

For images labelled for Gle1, isolated and spur pixels were removed from the image after binarization, along with areas of the binary image where the mean intensity in the corresponding grayscale image was smaller than 10% of the maximum grayscale value. Next the binary image was morphologically closed with a 5-pixel radius disk structuring element, followed by a dilation with a 4-pixel radius disk structuring element.

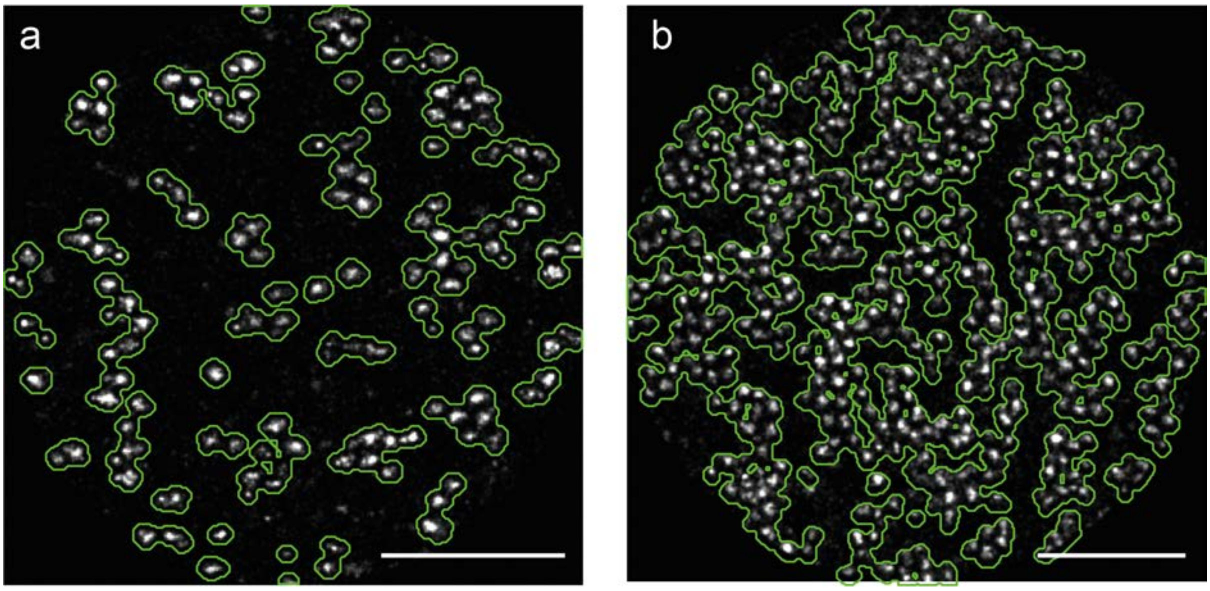


**Supplementary Figure 13: Binarization of super-resolved images labelled for Gle1**: Green lines indicate the boundaries between black and white pixels in binarized super-resolved images and are super-imposed on the original greyscale image for comparison purposes. Scale bars: 1 µm (**a**) A binarized super-resolved image of NPCs labelled for Gle1 in a follicle cell in a *Drosophila* stage 8 egg chamber with 19% NPC coverage of the nuclear membrane. (**b**) A binarized super-resolved image of NPCs labelled for Gle1 in a nurse cell in a *Drosophila* stage 4 egg chamber with 53% NPC coverage of the nuclear membrane.

**Estimating mean cluster distance**

The binary images were Euclidean distance transformed (**Supplementary Figure 14B**). To select only the pixel values along the ridges, whose values correspond to the half-distance between clusters, a morphological thinning operation was performed to produce a skeleton of the binary image (**Supplementary Figure 14C**). A round mask was also applied to exclude edge and peripheral pixels (**Supplementary Figure 14D**). The mean of the grayscale pixel values in the distance transformed image traced by the skeleton was then multiplied by 2 to obtain an estimate on the mean cluster distance in pixels (1 pixel corresponds to 10 nm).

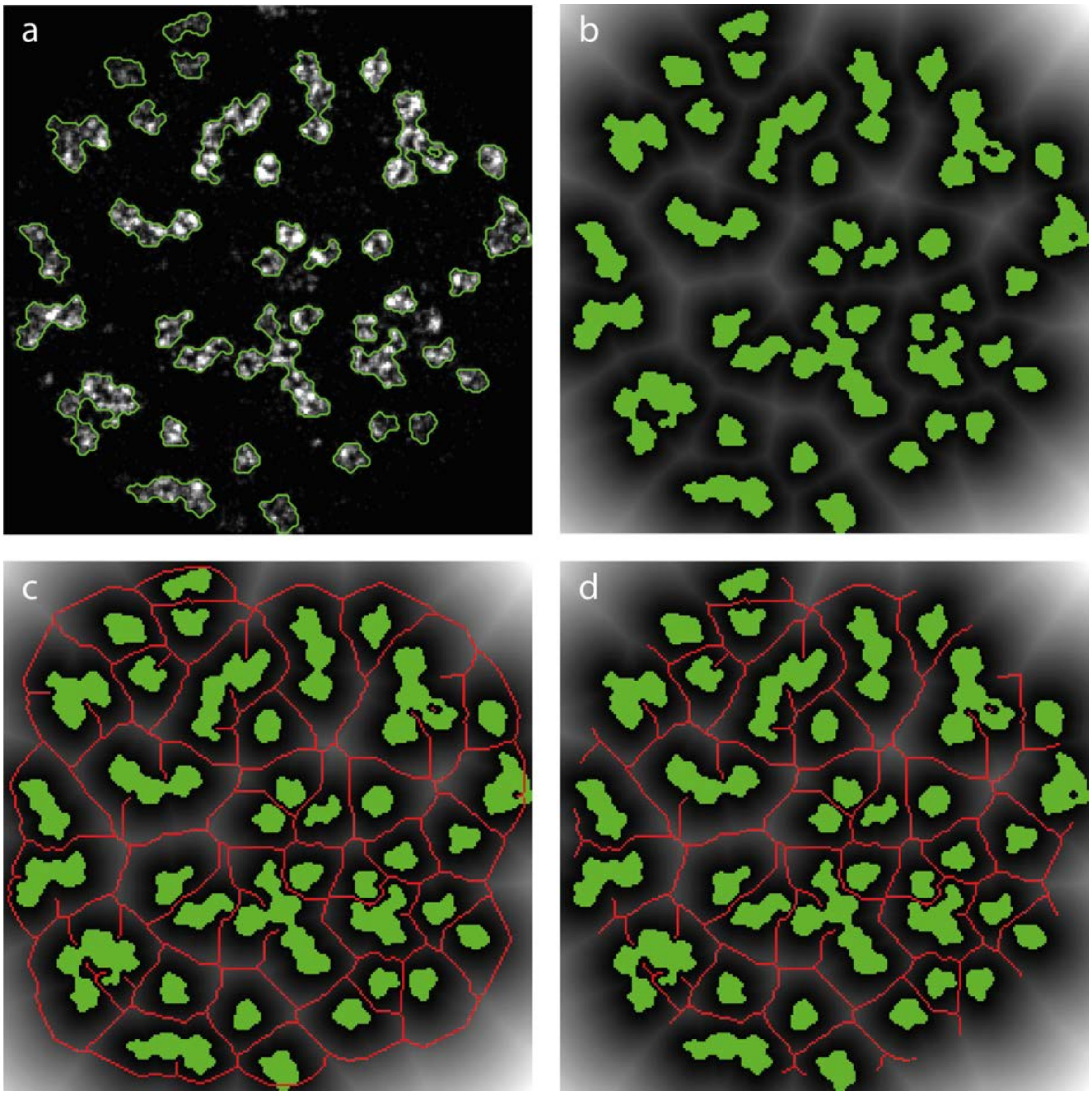

**Supplementary Figure 15: Mean NPC cluster distance estimation:** (**a**) A binarized super-resolved image of NPCs in a follicle cell in a *Drosophila* stage 8 egg chamber. Green lines trace the boundaries between white and black pixels and are superimposed on the original grayscale image. (**b**) Euclidean distance transform of the binary image shown in (a). The green areas indicate foreground pixels of the binary image in (a) and are superimposed on the grayscale distance transform image for convenience. The distance between the NPC clusters is estimated by the values of the distance transform image along the white ridges that run between the NPC clusters. (**c**) A skeleton produced by morphologically thinning the binary image is superimposed as a red trace on the distance transformed image. (**d**) The skeleton (red trace) after applying a round mask to exclude edge and peripheral pixels. The skeleton traces along the ridges in the distance transform image and provides a convenient way to select the pixels whose value corresponds to the distance between NPC clusters.

#### Estimating coverage

For the purpose of this work a binary image of a single NPC should appear as a disc with a radius of 5 pixels (50nm). However, the image binarization tends to assign a larger area to NPC clusters than is occupied by them (**Supplementary Figure 16**). To correct for this effect, we first analysed single NPCs and estimated the effective radius assigned by the binarization process. For data collected in the line scanning setup, a circular disk with an effective radius of 7.8 pixels was found to represent the area assigned to a single NPC. Indicating that the error introduced in the coverage by the binarization process is that of a circular ring with an inner ring radius 2.8 pixel smaller than its outer radius. The correction on the effective radius for the data collected in the TIRF system was 3.6 pixels.

For more complicated geometries associated with clusters of several NPCs and non-circular shapes, we calculated the radius that would correspond to a disk with the same area as the cluster under consideration. We apply the appropriate correction on this radius and the area calculated using this corrected radius was used for further data analysis.

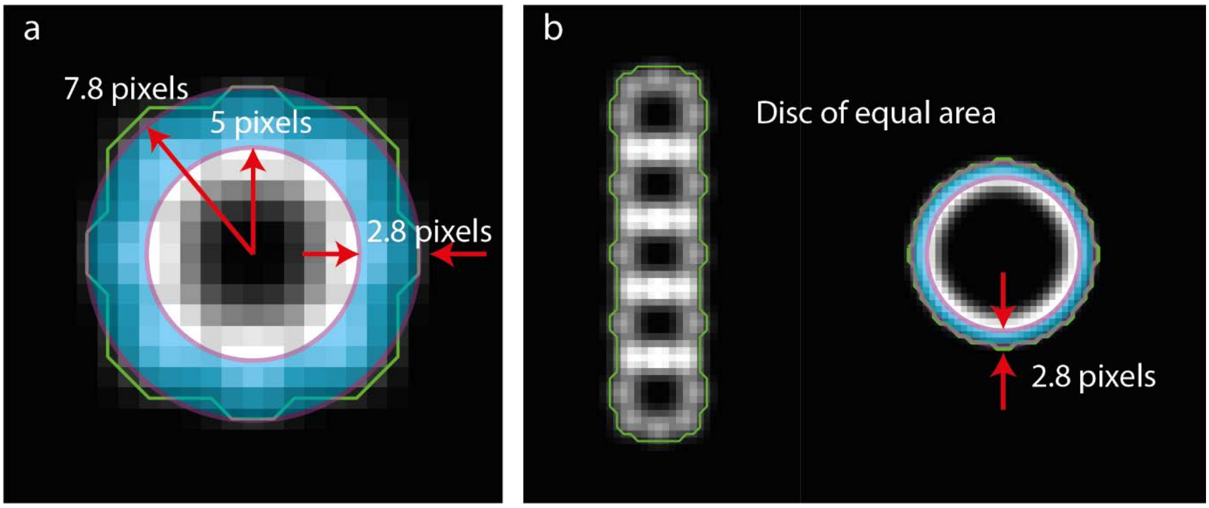

**Supplementary Figure 16: Estimating the coverage:** (**a**) Simulated image of a single NPC. The green trace indicates the area occupied by a single NPC after the binarization process. This area is approximated by a disc with an effective radius of 7.8 pixels. To obtain the actual area occupied by the NPC ,the value of the effective radius is reduced by 2.8 pixels and the area of a disc with 5-pixel radius is used. (**b**) Simulated image of a linear arrangement of 5 NPCs. The green trace indicates the area occupied by this NPC cluster after binarization. To estimate the actual area, we first calculate the effective radius of a disk with equal area as the one produced by the binarization process of this linear NPC cluster. The effective radius of this disc is reduced by 2.8 pixels, and the area of a disc with this reduced radius is used to obtain an estimate on the actual area occupied by the linear NPC cluster.

#### Simulation of NPC super-resolved images

To obtain a baseline for comparing the NPC cluster distribution between different tissues, we generated simulated NPC images by randomly distributing NPCs, under the condition that NPCs cannot overlap, and varied the number of NPCs to achieve coverages from 3-60%. For each coverage condition, datasets with a labelling efficiency ranging from 20-100% were generated. To factor in variability associated with the random distribution of NPCs, 10 repeats for each coverage and labelling condition were performed. The simulated images were morphologically analysed in an identical way to the super-resolved images obtained from the different tissues.

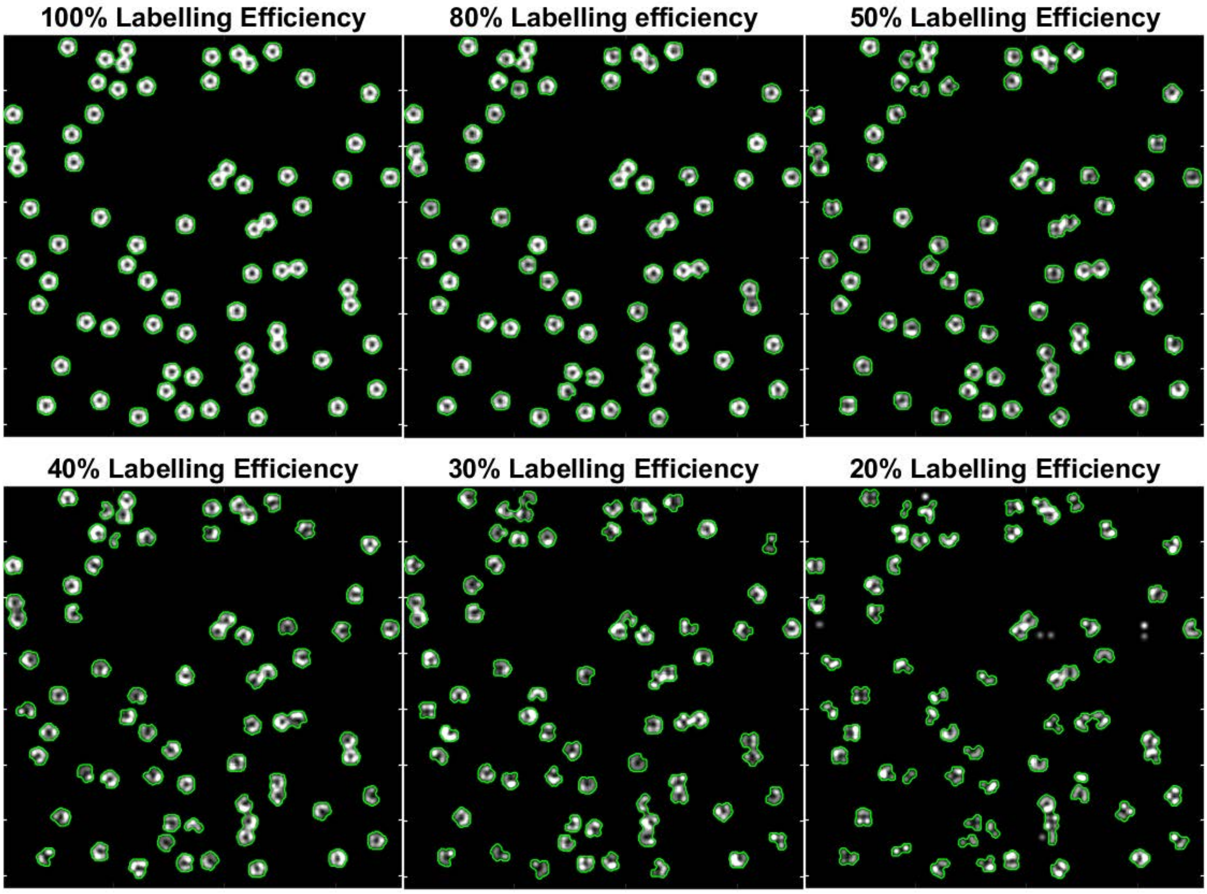

**Supplementary Figure 17: Overview of simulated NPC datasets with 6% coverage and different levels of labelling efficiency.**

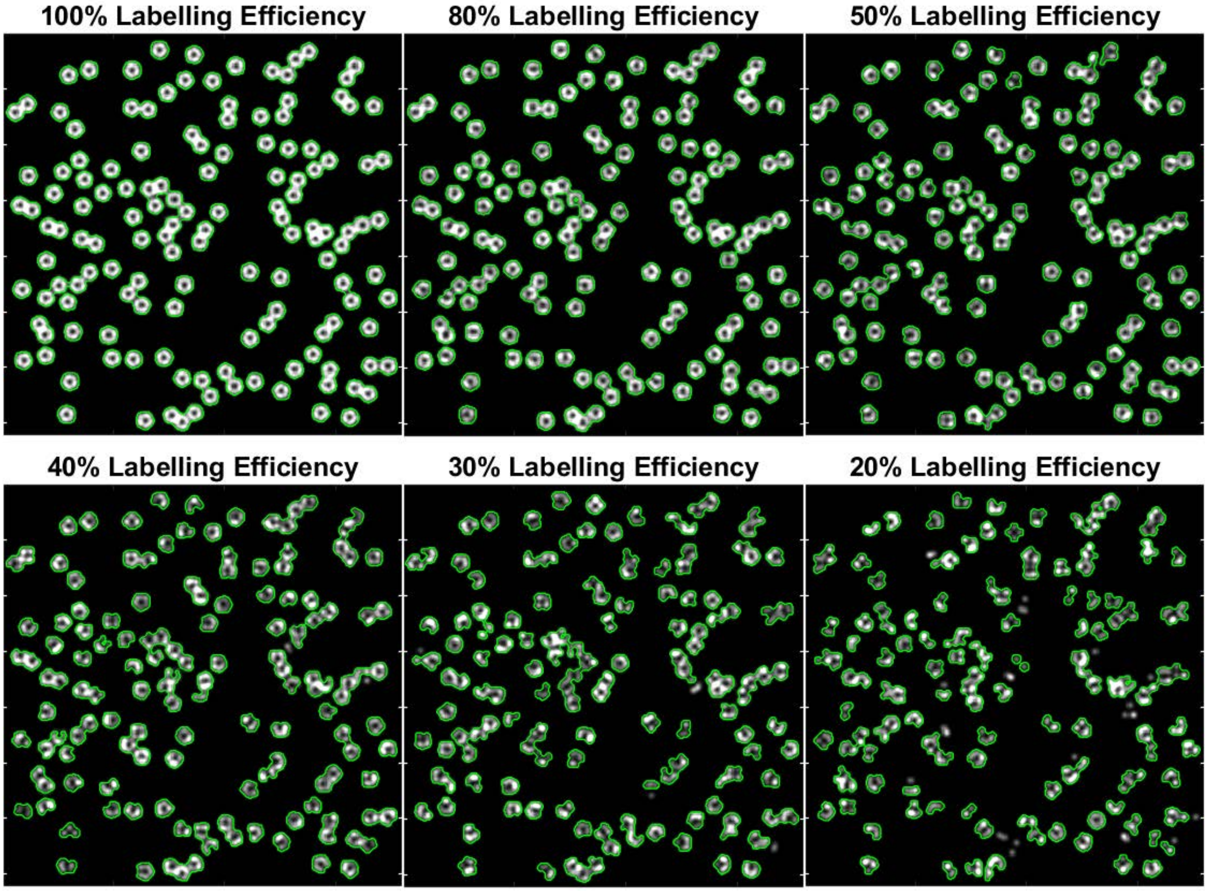


**Supplementary Figure 18: Overview of simulated NPC datasets with 12% coverage and different levels of labelling efficiency**

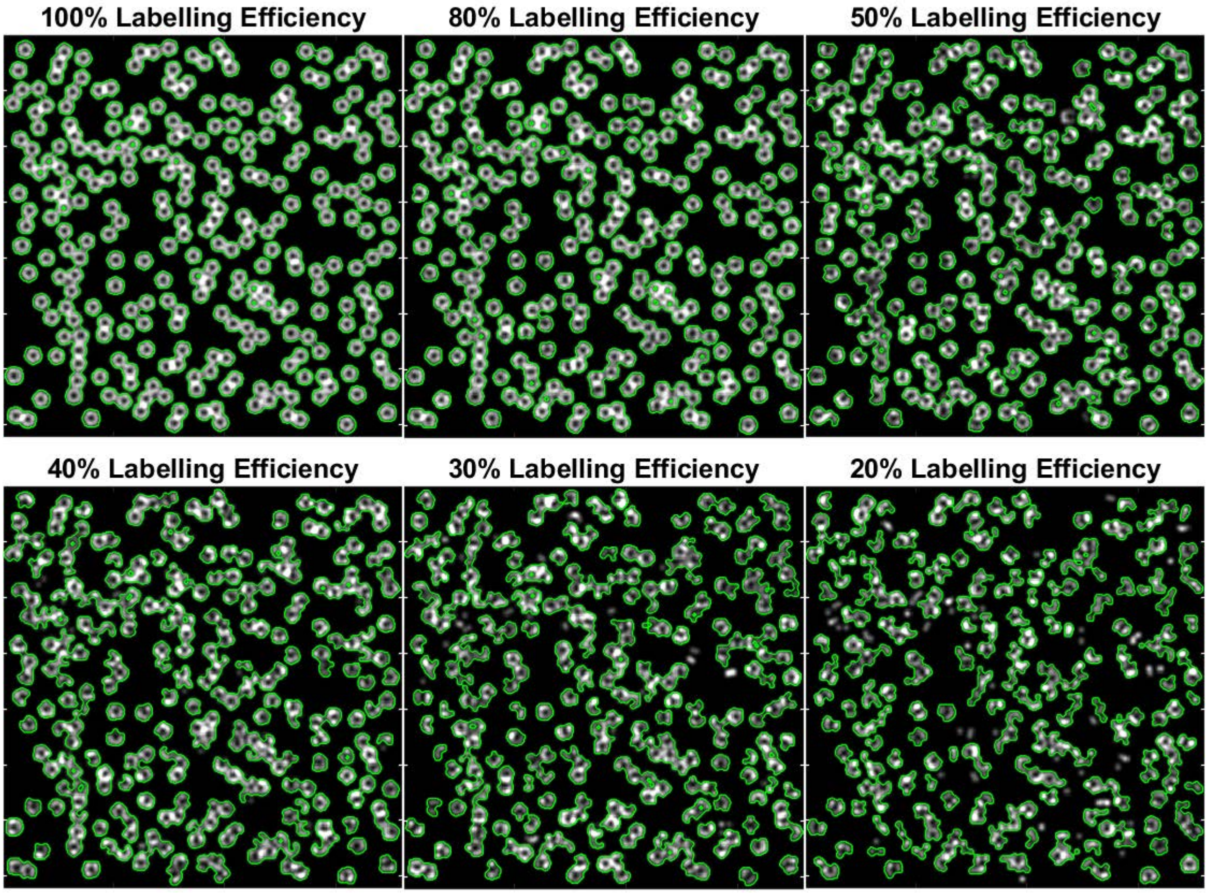


**Supplementary Figure 19: Overview of simulated NPC datasets with 24% coverage and different levels of labelling efficiency**

The mean NPC-cluster distance versus coverage for the different labelling conditions and corresponding repeats is shown in **Supplementary Figure 20**. Based on these data an expected trend is drawn by fitting the 50% label efficiency data with a power equation ($a * x^b$). Upper confidence intervals are estimated by selecting only the highest NPC-cluster distance value at each coverage value and fitting those values with the same power equation. Lower confidence intervals are calculated in a similar way, but by selecting the corresponding lowest NPC-cluster distance at each coverage value.

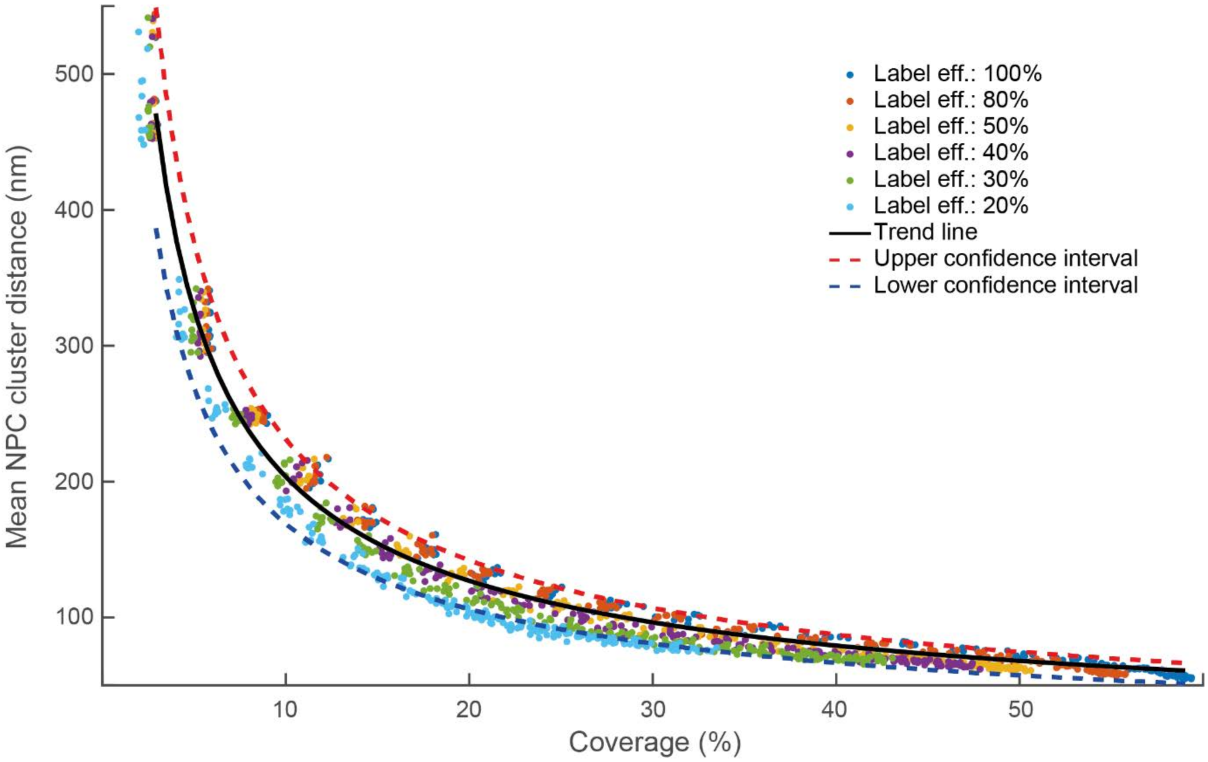


**Supplementary Figure 20: Mean NPC-cluster distance vs Coverage.**

#### Optimizing Imaging Parameters

The excitation laser intensity is a critical parameter with a profound effect on the quality of super-resolved images [5]. Within the context of line-scanning DNA-PAINT we found it important to optimize laser intensity to increase the number of photons per localization without negatively impacting the labelling efficiency due to photodamage. Although DNA-PAINT is resistant to photobleaching due to the abundance of free-floating dyes attached to imager strands, docking strands are not exchangeable and are susceptible to photodamage[6]. To avoid depleting docking strands due to photodamage we monitored the number of localizations per frame. For Atto 643 and Cy3B, we found that a diffraction limited excitation line with less than 2 mW power maintained a reasonably constant number of localizations throughout the course of imaging as shown in **Supplementary Figure 21a-b**.

Under this excitation condition we then varied the slit width to obtain an optimum slit width of 16 lines (**Supplementary Figure 21c**)

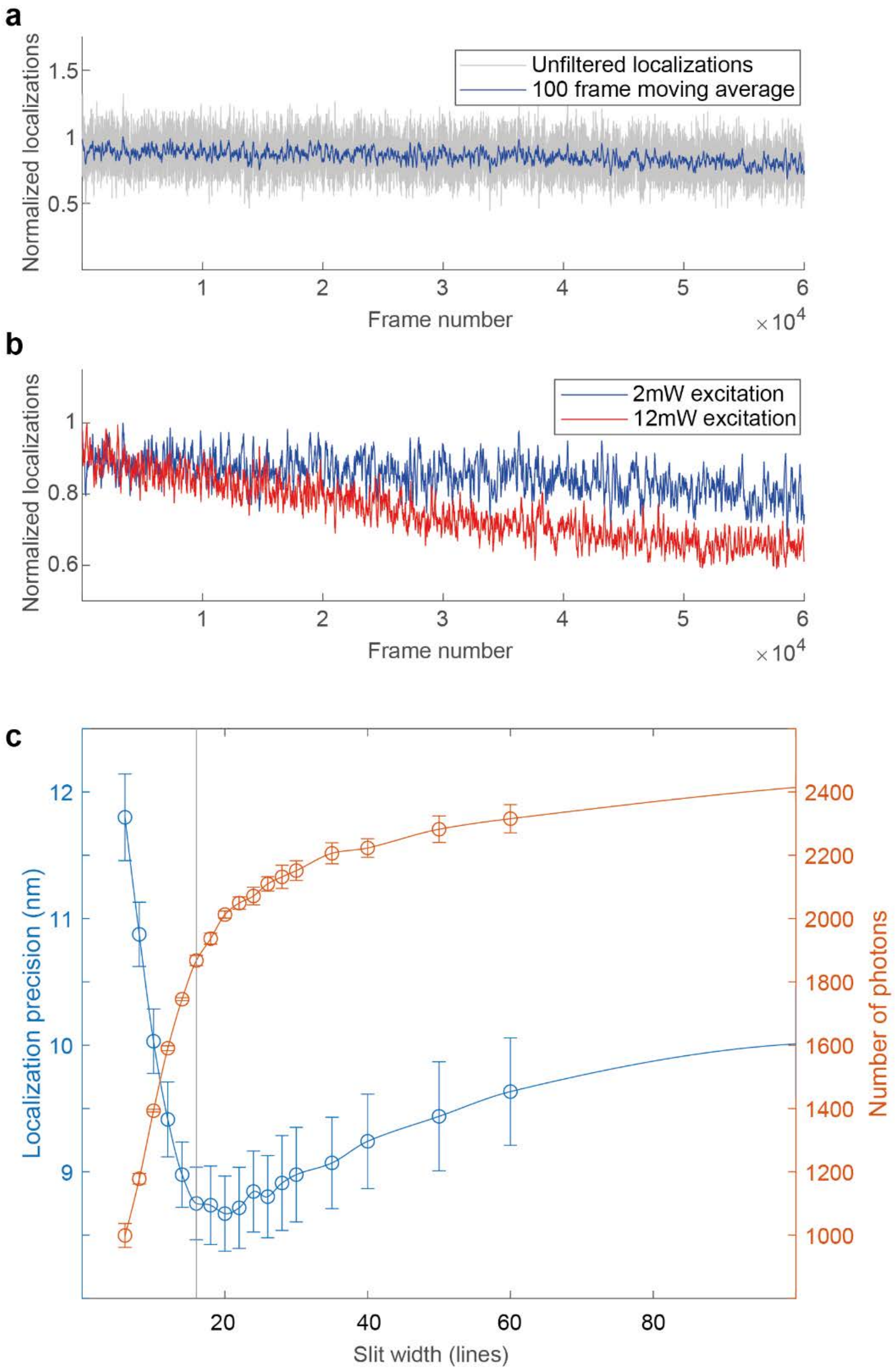

**Supplementary Figure 21: Optimization of imaging parameters:** (**a**) Number of localizations as a function of frame number. Gray trace indicates the raw number of localizations and the blue trace the moving average over 100 frames under 2mW excitation and 4Hz framerate. **(b)** 100-frame moving average of localizations as a function of frame number under 2mW excitation (blue trace) and 12mW excitation (red trace) recorded at 4Hz framerate. (**c**) Mean localization precision (blue trace) and mean number of photons (red trace) as a function of slit width of 500 frames under 20% excitation recorder at 4Hz frame rate in the presence of 1nM Atto643 imager strand solution. Error bars indicate mean ± s.d. for 3 independent experiments.

#### Selecting Line ROIs and Binning Localization Data

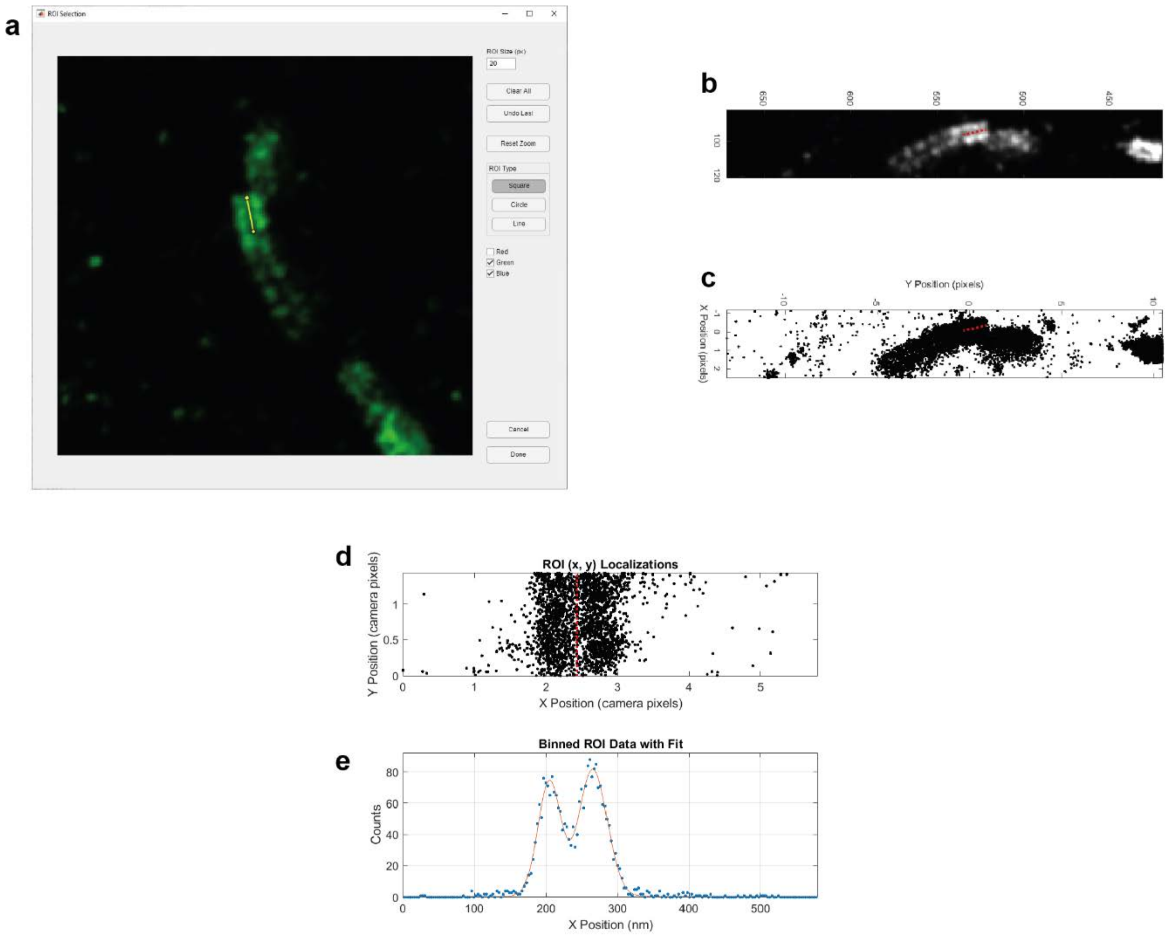


**Supplemental Figure 22. Localization selection, rotation, and binning. a** A simple ROI selection interface allows the user to draw a line, or lines, on a reconstructed image. The image show here is the same as the **Main Text Figure 2**, specifically the NPC profile used in panels (2c) and (2d). The user drawn line is shown in yellow. **b** and **c** After one or more lines have been draw on the reconstructed image (red dashed line), the underlying (x, y) localizations around the line are selected to be used in the next step. **d** The angle of the user drawn line is computed and the localizations are rotated by an equal angle such that the original user drawn line (red dashed line) would be parallel to the vertical axis. **e** (x, y) localizations are binned along the y-axis to produce a profile that is the equivalent to having summed the localizations along the original user drawn line.

Within the main text, **Figures 2** and **3** include cross sections of image features. More specifically, in **Figure 2**, panels **c** and **d** we show the results of measuring the distance between the cytoplasmic and nucleoplasmic rings of a nuclear pore complex (NPC) in profile view and in **Figure 3**, panels **a''**, **b''**, and **c''** we show the results of measuring the diameter of small, straight, filament structures. We also report the mean value for measurements of multiple NPCs ring separation, filament structures seen in our actin images and the distance between adjacent plasma membranes observed on the apical surface of our actin images. In all instances, rather than taking a simple line profile with pixel intensities from the reconstructed image we have used the underlying (x, y) localization data. To accomplish this, we have developed several custom MatLab scripts. Here we present a short overview of the process.

(x, y) localization data is loaded into MatLab and used to generate a reconstructed image. This reconstructed image is displayed in a custom made ROI selection window (app) inside the MatLab environment (**Supplementary Figure 22a**). The user can select square, circular, or line type ROIs. In all cases, the line tool was employed. The user may zoom and pan the image while drawing lines on the target features. After the user has finished drawing ROI lines, the ROI selection tool is closed and the line start and end points are converted from image pixel space into (x, y) localization space. The (x, y) localizations within a user set distance from the line ROI (typically ±3 camera pixels, or ±300 nm) are copied into a corresponding ROI variable (**Supplementary Figure 22b and c**). Based on that start and end points of the line, the angle and mid-point of the line are computed. The (x, y) midpoint is subtracted from the (x, y) localization coordinates such that the centre of the line ROI now resides at (0, 0). The (x, y) localizations are next converted into polar coordinates and the line ROI's angle is added to the θ component of the (r, θ) localizations positions. The localization coordinates are transformed back into cartesian (x, y) coordinates. When plotted (**Supplementary Figure 22d**), the feature over which the original user drawn line was placed is now parallel to the vertical axis. These rotated (x, y) coordinates are binned to produce a feature profile that is equivalent to having summed the localization along the line ROI and may be fit to determine the width of the feature or, as shown in **Supplementary Figure 22e**, the distance between to features.

#### Estimating the Microscope's Resolution

As is the case for any localization based microscope, the resolution is depended on several factors including the number of collected photons per blink per camera frame, the number of background photons, sample drift, and the labelling density of the structure being imaged. Although computational methods have been reported for determining the resolution of reconstructed SMLM images, here we simply report the results of fitting image features as described in the **Selecting Line**

**ROIs and Binning Localization Data** section. Feature size within SMLM images give a native estimate of what resolution may be possible.

#### NPC Ring Profile Fitting

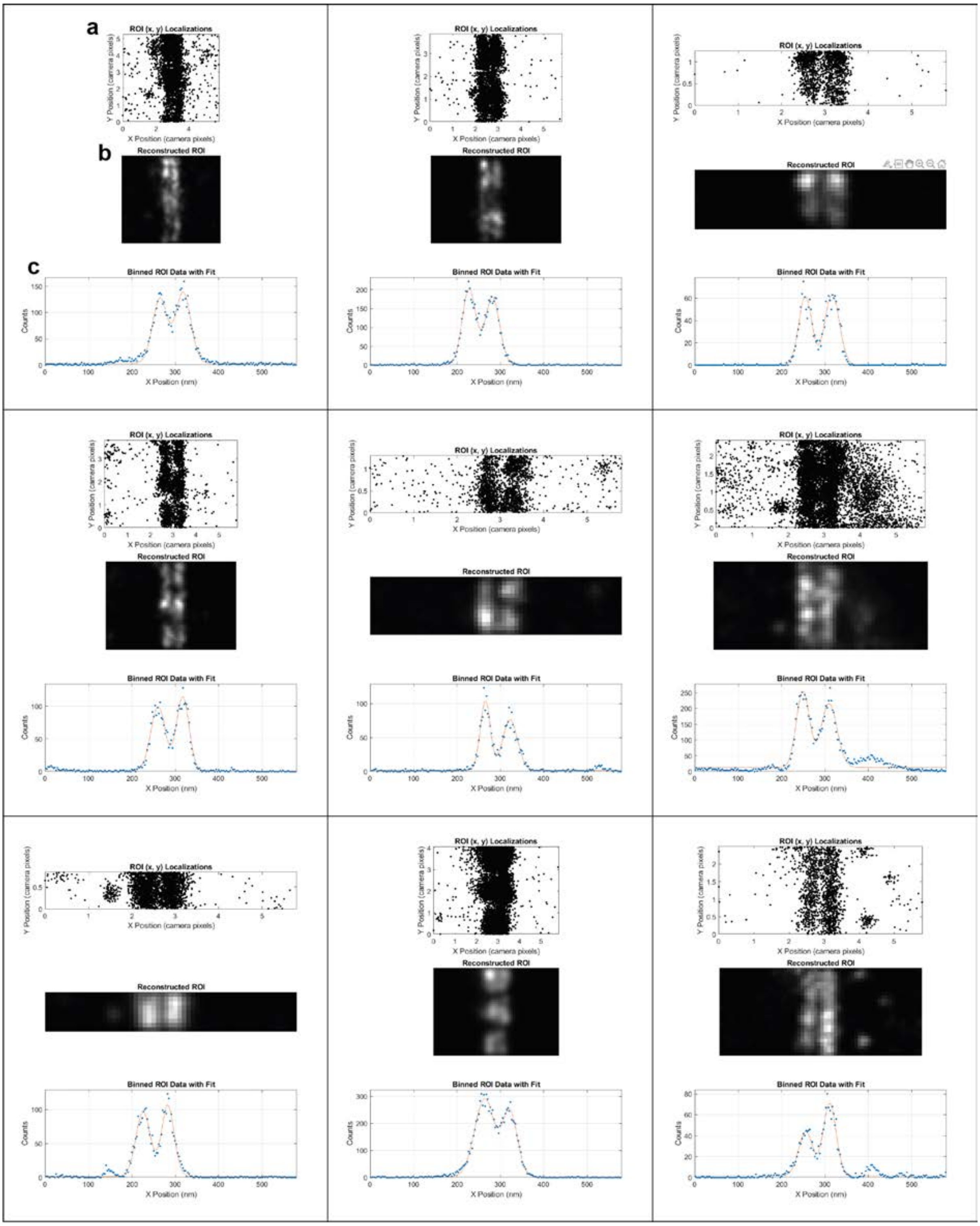


**Supplemental Figure 23. Typical NPC cross-sectional views binned and fit with two Gaussians.** A line is drawn on a reconstructed SMLM image and the localizations lying along the line, and within a certain distance of the line, are segmented out of the (x, y) localization dataset and rotated such that the NPC profile is parallel to the vertical axis as shown in **a**. **b** The segmented (x, y) localizations may be reconstructed into a typical SMLM image. **c** Localization are binned (blue data points) and fit (solid red line) with two gaussians to determine the distance between the cytoplasmic and nucleoplasmic rings.

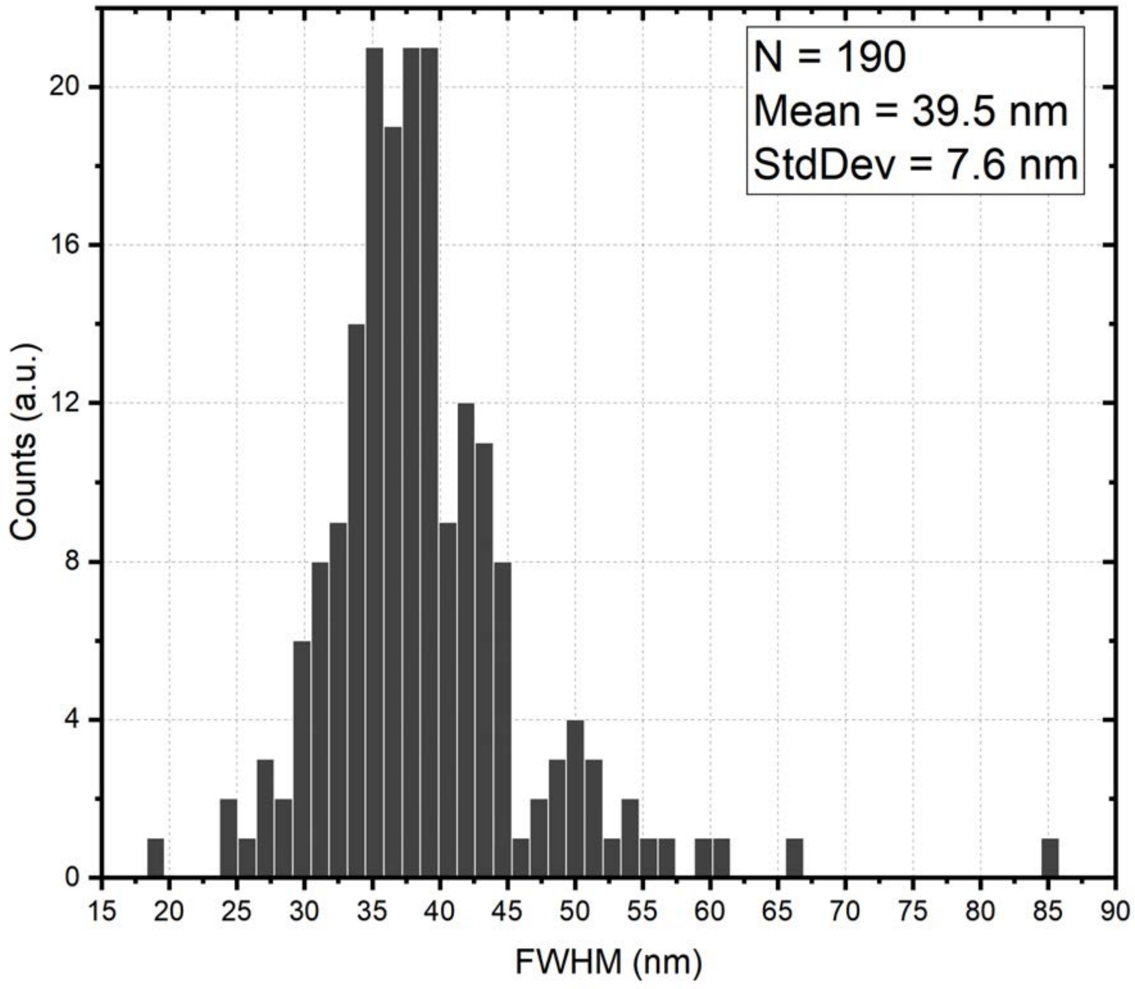


**Supplemental Figure 24. Distribution of fitted NPC cross-sectional ring widths.** Following from the **Supplemental Methods** and **Supplemental Figure 23**, 190 NPC cross-sections were hand selected. The (x, y) localization coordinates for each NPC cross-section were rotated, binned and fit with a double Gaussian function to determine the mean size for a single ring in profile resulting in a mean value of 39.5 ± 7.6 nm (mean ± standard deviation, N = 190).

Imaging nuclear pore complexes on the nuclear membrane of nurse cells necessitates a focal plane deep inside an egg chamber. As seen in **Main Text Figure 2b**, at this focal depth, imaging often catches a nuclear envelop cross section within the image and, subsequently, cross-sectional views of NPCs (Nup160, boxed region **Main Text Figure 2c**). These cross-sectional views afford an opportunity to (1) determine the separation between an NPC's cytoplasmic and nucleoplasmic rings (**see Supplemental Figure 4**) and (2) determine the fitted width of a single NPC ring in cross-section as a metric for resolution. As such, 190 Nup160 NPC cross-sections were hand selected and processed as described in the **Selecting Line ROIs and Binning Localization Data** section above. Representative ROIs and fits of Nup160 NPC cross sections are presented in **Supplementary Figure 23**. The mean full width at half maximum (FWHM) of a single NPC ring in cross-section was measured as **39.5 ± 7.6** nm (mean ± standard deviation, N = 190). The distribution of measured distances between the rings is shown in **Supplementary Figure 24.** As a single Nup160 NPC ring in cross section should be below the resolving power of the system described here, this should give a realistic idea of how large nanoscale structures will appear on this system under these imaging conditions (see **Supplementary Table 3** for typical imaging conditions).

**Actin Filament Feature Fitting**

Single actin and intermediate actin filaments are known to have diameters in the single digit nanometre range. Thus, imaging actin offers another real-world opportunity to measure the resolving power of the system described here. As such, actin was imaged as described within the main text and **Supplementary Table 3**. As previously described within the Supplementary Methods, small features within the actin images presented in **Main Text Figure 2** were annotated by hand, the localization were binned, and the effective profile was fit with a Gaussian to determine the FWHM. Representative ROIs and fits of actin filaments can be seen in **Supplementary Figure 6**. In total **21** filaments were selected and fit, **11** from the basal actin image (**Main Text Figure 2a**) and **10** from the apical actin images (**Main Text Figure 2b and c)**. The measured FWHM of the small actin features (presumably filaments) selected in these images is **30.3 ± 6.67** nm (mean ± standard deviation, N = 21). The distribution of fitted FWHMs is shown in **Supplementary Figure 7.** Features in the basal actin image were found to have an average size of **30.4 ± 8.42** nm (mean ± standard deviation, N = 11) vs. apical actin with an average size of **30.2 ± 4.48** nm (mean ± standard deviation, N = 10) suggesting no major reduction in resolving power between basal and apical surfaces under these imaging conditions (see **Supplementary Table 3** for typical imaging conditions). As actin filaments should be below the resolving power of the system described here, this should give a realistic idea of how large nanoscale structures will appear on this system under these imaging conditions (see **Supplementary Table 3** for typical imaging conditions).
